# OrthoSeq: A Design Workflow for Thermodynamically Orthogonal DNA Sequence-Pair Libraries

**DOI:** 10.64898/2026.08.01.742265

**Authors:** Florian Katzmeier, Matthew Aquilina, William M. Shih

**Affiliations:** Department of Cancer Biology, Dana-Farber Cancer Institute, Boston, MA 02215, USA; Department of Biological Chemistry and Molecular Pharmacology, Harvard Medical School, Boston, MA 02115, USA; Wyss Institute for Biologically Inspired Engineering, Harvard University, Boston, MA 02215, USA

## Abstract

Programmable DNA hybridization underlies many technologies in DNA nanotechnology, fluorescence imaging, and synthetic DNA sequence assembly. A common design challenge is to generate large sequence libraries in which each strand binds strongly to its intended partner while avoiding cross-hybridization and self-folding. Here, we introduce OrthoSeq, a workflow for designing thermodynamically orthogonal DNA sequence-pair libraries under user-defined experimental conditions. OrthoSeq uses NUPACK to evaluate intended binding, cross-hybridization, and self-folding. Within OrthoSeq, candidate sequence pairs form vertices in a conflict graph, while pairwise cross-hybridization conflicts define the edges. Library selection is then formulated as an independent-set problem and addressed using search strategies tailored to the computational regime considered here, in which thermodynamic evaluations dominate the computational cost. In benchmark comparisons, these strategies identify larger sequence-pair libraries than the commonly employed sequential candidate-addition strategy under the same thermodynamic constraints and computational budget. We further show that sequence-level barcode libraries can serve as candidate pools for thermodynamic refinement with OrthoSeq. To support practical use, OrthoSeq provides a graphical user interface that implements the complete workflow. Altogether, OrthoSeq provides an application-agnostic framework for designing DNA sequence-pair libraries under explicit thermodynamic constraints.

## Introduction

In many DNA-based technologies, single-stranded DNA sequences act as programmable binding sites that determine which molecular or nanoscale components bind to one another. This principle is used to assemble DNA origami units into finite structures (*1* –*3*), organize DNA nanostructures into periodic crystals (*4*, *5*), and control interactions between DNA-coated colloids and nanoparticles (*6*). Short sequences also act as toeholds in dynamic DNA nanotechnology and synthetic reaction networks (*7*), and as local binding rules in algorithmic molecular self-assembly (*8*). Sequence-programmed hybridization is also central to multiplexed fluorescence imaging. In MERFISH (*9*) and seqFISH (*10*), repeated rounds of hybridization to readout probes and imaging allow many transcripts to be identified and localized in fixed cells or tissues. In DNA-PAINT (*11*) and Exchange-PAINT (*12*), transient binding of fluorescent imager strands to docking sequences enables super-resolution imaging and multiplexed target readout. More recently, Sidewinder combined orthogonal auxiliary sequence pairs with DNA three-way junctions to build long, complex sequences from short oligonucleotides, separating the assembly-guiding information from the final product (*13*). Across all these applications, the same design problem arises: sequence libraries must contain many sequences that bind with their intended partners while minimizing cross-hybridization with other sequences in the library.

Existing sequence design methods can be broadly divided into two classes. The first optimizes a sequence set of predetermined size to promote intended binding while reducing cross-hybridization. For this optimization, one or more sequence-quality scores are defined, and candidate sequences are improved through stochastic changes such as mutations, crossovers, or local sequence updates (*8*, *14*). Because these scores combine the contributions of many interactions, a favorable overall score can still coexist with one or more strong unintended interactions. This approach therefore produces a library of the requested size, but it does not necessarily ensure that every unintended interaction remains below a specified cross-hybridization limit. It is also well suited for designing complex DNA structures and reaction systems with interaction patterns beyond simple pairwise binding, as exemplified by the NUPACK multistate design tool (*15*).

The second class instead seeks to identify the largest possible library of sequence pairs for which intended binding is sufficiently strong and every unintended interaction remains below a predefined cross-hybridization limit. With fixed interaction limits, sequence selection can be formulated as a graph-theoretic optimization problem. Candidate sequence pairs are represented as vertices, and an edge connects two vertices if the corresponding sequence pairs cannot coexist without violating a cross-hybridization limit. The goal is then to identify a large independent set representing mutually compatible sequence pairs (*16* –*18*). This approach ensures that all selected sequence pairs satisfy the specified constraints, but the resulting library size is not known in advance.

Independent of the design objective, sequence design methods also differ in how they quantify sequence interactions and cross-hybridization. Many existing library-selection methods use computationally efficient sequence-level criteria, including sequence complementarity, simplified nearest-neighbor-based scores, Hamming-distance constraints, or sequence-symmetry minimization (*16* –*19*). These criteria allow large candidate pools to be evaluated efficiently, but they do not directly impose thermodynamic binding limits under specified experimental conditions. We encountered this limitation while expanding sequence libraries for micron-scale crisscross DNA origami (*3*), which motivated the more general workflow developed here.

Here, we introduce OrthoSeq, a workflow that combines the library-selection objective of the second class with explicit NUPACK-based thermodynamic criteria for intended binding, cross-hybridization, and secondary-structure formation under user-defined experimental conditions. We formulate sequence selection as an independent-set problem in a conflict graph and use a graph-aware search strategy (*20*) as an alternative to the commonly employed sequential candidate-addition strategy (*16*, *17*, *21*). For sequences up to 7 nucleotides, the full conflict graph can be constructed, and graph-aware search finds larger sequence libraries than sequential candidate addition under the same thermodynamic constraints. NUPACK-based cross-hybridization evaluation is computationally expensive, and exhaustive pairwise evaluation becomes limiting for longer sequences. We therefore introduce a hybrid search strategy that combines graph-aware search with computationally efficient sequential candidate addition. Under a fixed computational budget, this hybrid strategy consistently finds larger sequence libraries than sequential candidate addition alone for sequence lengths ranging from 8 to 25 nucleotides. Together, these strategies cover the full tested sequence-length range by exploiting the complete conflict graph when feasible while remaining computationally practical when exhaustive graph construction is prohibitive.

To compare sequence-level heuristics with thermodynamic criteria for orthogonality, we evaluate OrthoSeq against seqwalk, a recently developed method for generating large barcode libraries through sequence-symmetry minimization (*19*). Unlike the two optimization classes described above, seqwalk constructs libraries directly using a sequence-level heuristic rather than optimizing a predefined design or selecting a subset under fixed interaction constraints, enabling computationally efficient generation of large libraries. This comparison highlights that satisfying sequence-level orthogonality criteria does not necessarily imply thermodynamic orthogonality. We also show that seqwalk-generated barcode pools can be refined by OrthoSeq when predictable hybridization behavior or additional user-defined constraints are required.

Finally, we implemented the complete OrthoSeq workflow in an open-source Python package that supports both scripting and a graphical user interface for custom library generation under user-defined experimental conditions.

## Materials and Methods

### Thermodynamic criteria for sequence selection

In this work, a sequence pair consists of a DNA strand and its intended binding partner. Each sequence consists of optional 5*^′^* and 3*^′^* extensions and a core binding domain of defined length. The intended binding partner is derived from the reverse complement of the core binding domain, as illustrated in Figure 1A. We seek to identify large sets of such pairs without appreciable cross-hybridization. Specifically, each strand should bind strongly to its intended partner (on-target interaction), but weakly to all other strands in the set (off-target interactions).

**Figure 1:**
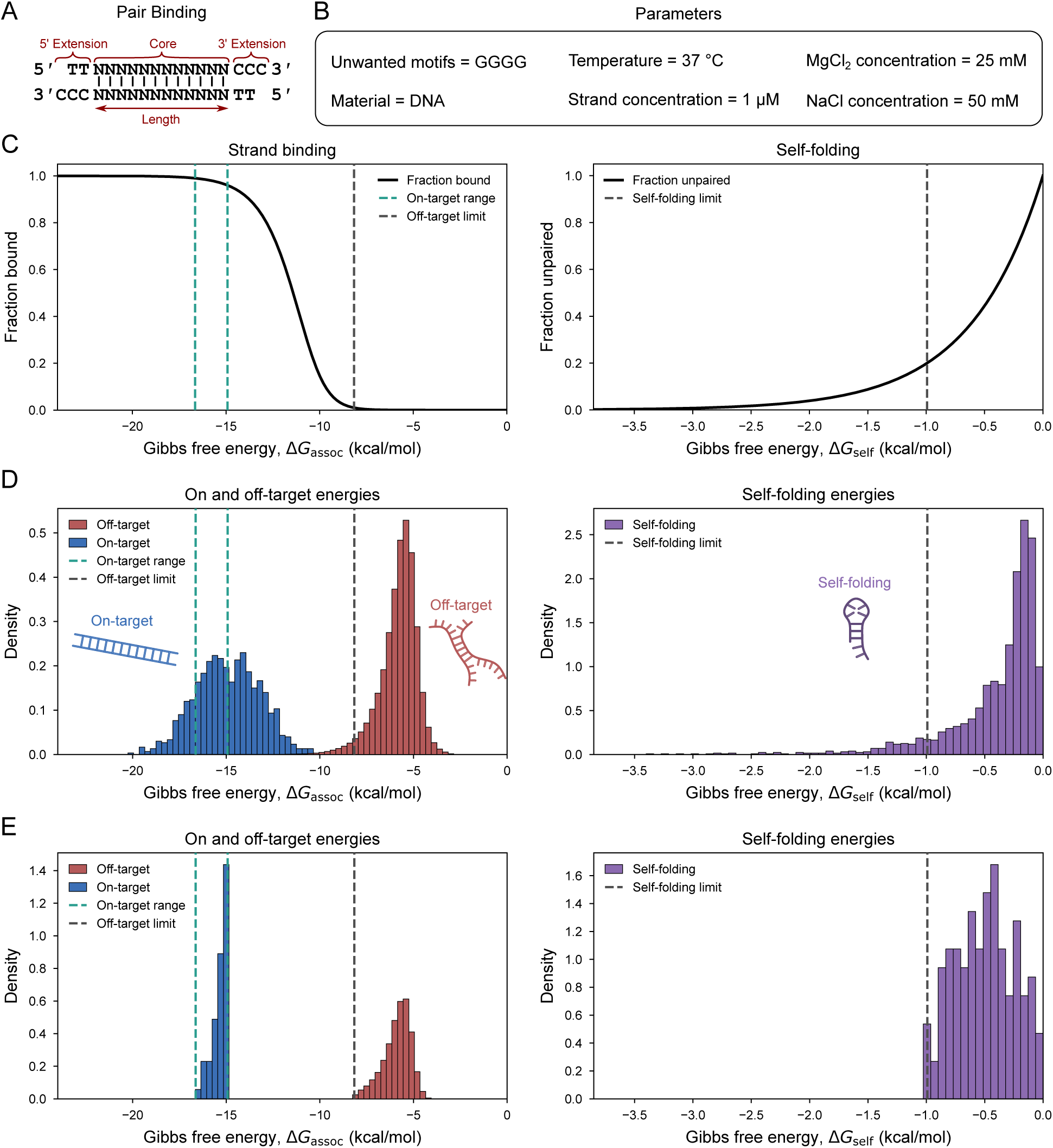
Thermodynamic problem formulation for orthogonal sequence-pair selection. **(A)** Definition of a sequence pair. Each sequence consists of optional 5*^′^* and 3*^′^*extensions and a core binding domain of definable length. Binding between the core binding domain and its intended binding partner is indicated. **(B)** Physical parameters used for generating the plots shown in panels C, D, and E. **(C)** Relationship between fraction bound and association free energy, Δ*G*_assoc_ (left), and between fraction unpaired and self-folding free energy, Δ*G*_self_ (right). Vertical lines indicate example limits used to define allowed energy ranges. **(D)** Energy distributions for 1000 random 12-mer sequence pairs obtained using NUPACK with the physical parameters shown in panel B. Vertical lines indicate the same example limits as in panel C. **(E)** Energy distributions of a representative orthogonal set obtained using our sequence-selection algorithm. The distributions fall within the allowed energy ranges defined in panel C.

These criteria can be quantified using a representative two-strand equilibrium reaction of the form *A* + *B* ⇌ *AB*. We define strong binding as a large fraction of strands *A* and *B* being bound in the complex *AB*, whereas weak binding corresponds to only a small fraction being bound in the complex. In practical terms, intended on-target interactions must exceed a chosen limit on the fraction bound, whereas off-target interactions are required to remain below a chosen limit on the fraction bound. The fraction bound can be mathematically related to the association free energy, Δ*G*_assoc_, through the equilibrium constant of the isolated association reaction. Thus, on-target and off-target limits on the fraction bound can be directly translated into limits on the association free energy.

A second important criterion is that each individual strand in the set should have a weak tendency to form intramolecular secondary structures, hereafter referred to as self-folding. This criterion can similarly be quantified using a representative intramolecular equilibrium reaction of the form *U* ⇌ *S*, where *U* denotes the fully unpaired state and *S* represents the ensemble of self-folded states. We define weak self-folding as a large fraction of strands being in the fully unpaired state at equilibrium. In practical terms, this means that the fraction of strands in the fully unpaired state must exceed a chosen threshold. As for intermolecular binding, this threshold can be translated into a limit on the self-folding free energy, Δ*G*_self_. Calculation details for the free-energy thresholds, including an explicit treatment of homodimer formation, are provided in Section 1 of the Supporting Information.

The association and self-folding free energies were calculated with the NUPACK thermodynamic model using the DNA parameter set dna04.2 (*22*). NUPACK calculates ensemble free energies of complexes and individual strands from the corresponding partition functions, which we use to compute the association and self-folding free energies (*23*). For a two-strand association reaction *A* + *B* ⇌ *AB*, we define the association free energy as Δ*G*_assoc_ = *G_AB_* − *G_A_* − *G_B_*, where *G_A_*, *G_B_*, and *G_AB_* are the free energies of strand *A*, strand *B*, and the two-strand complex, respectively. The self-folding free energies are identical to the free energies of strands *A* and *B*, *G*_self*,A*_ = *G_A_* and *G*_self*,B*_ = *G_B_*.

The relationships between fraction bound and association free energy, Δ*G*_assoc_, and between fraction unpaired and self-folding free energy, Δ*G*_self_, are illustrated in Figure 1C for the parameters shown in Figure 1B, with vertical lines indicating example limits. The on-target binding criterion also includes a lower free-energy limit to avoid excessively strong strand binding.

In Figure 1D, we plot the distributions of the association free energy, Δ*G*_assoc_, for on-target and off-target interactions, together with the distribution of the self-folding free energy, Δ*G*_self_, for 12-mer sequence pairs. The same example limits as in Figure 1C are shown as vertical lines, indicating the allowed energy ranges for selecting an orthogonal set of sequence pairs. The desired outcome of sequence-pair selection is illustrated in Figure 1E, which shows the energy distributions of a representative orthogonal set obtained using the algorithm described below. The distributions fall within the same on-target, off-target, and self-folding limits indicated in Figure 1D.

### Conflict-graph interpretation

In this section, we describe how our thermodynamic criteria can be used to formulate the selection of orthogonally binding DNA sequence pairs as a maximum independent set problem in a conflict graph, as illustrated in Figure 2A.

**Figure 2:**
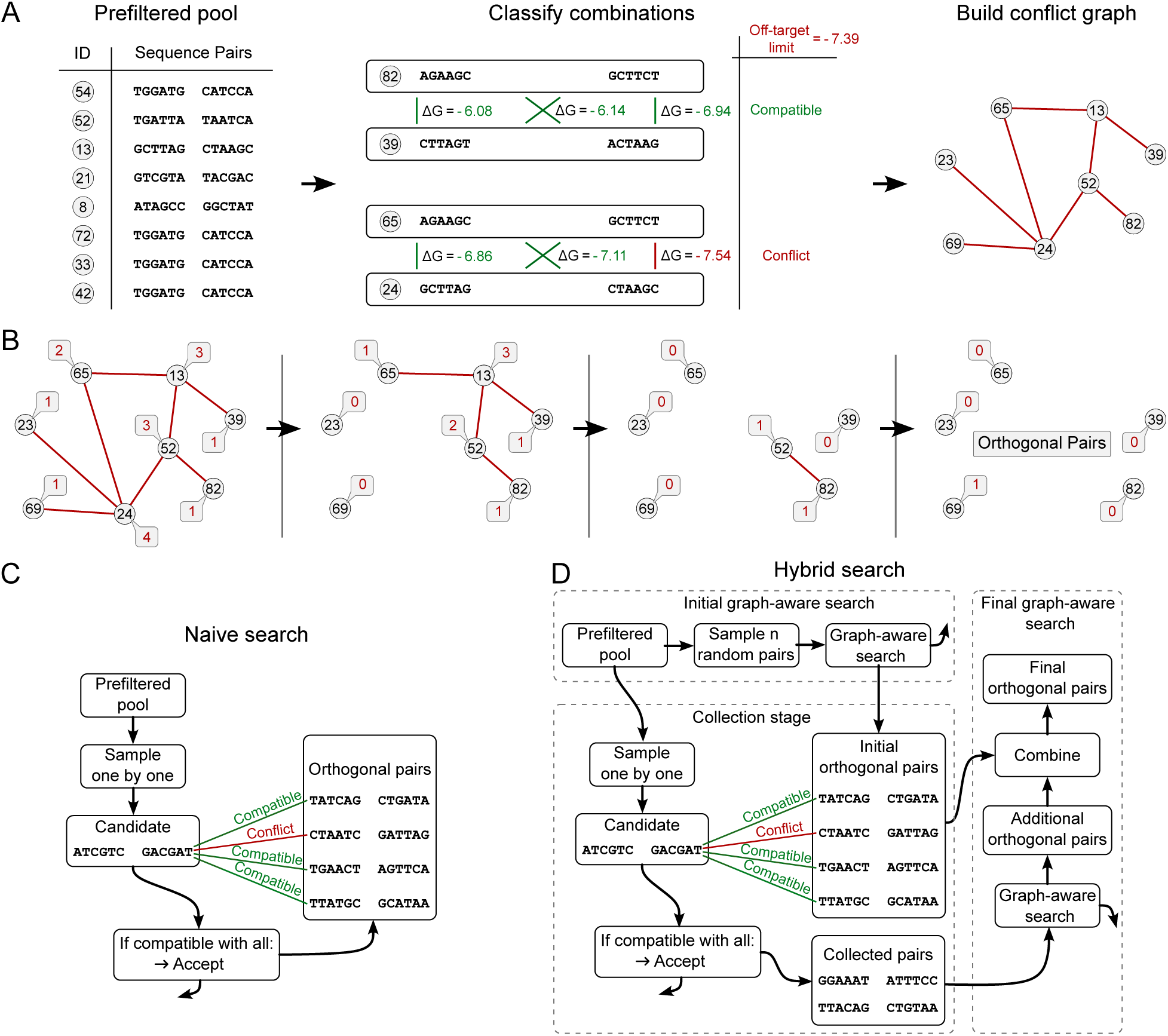
Search strategies for orthogonal sequence-pair selection. **(A)** Conflict-graph formulation. Left: sequence pairs form a prefiltered pool and are assigned ID numbers. Middle: two example combinations of sequence pairs are shown. Each combination has four possible off-target interactions, indicated by green lines. Free energies are shown for three selected interactions only and are reported in kcal/mol. If any interaction exceeds the chosen off-target limit, the combination is classified as conflicting. Right: resulting conflict graph, where vertices represent sequence pairs and edges indicate conflicts. **(B)** Graph-aware search. Each vertex is labeled with its number of edges. From left to right, vertices with the highest number of edges are eliminated iteratively, and the number of edges is then recounted for the remaining vertices. When no edges remain, the remaining vertices form an independent set. **(C)** Naive search. Sequence pairs are considered one by one and added to the independent set only if they have no off-target conflict with any sequence pair already in the set. **(D)** Hybrid search. The strategy proceeds in three stages, indicated by dashed boxes. First, an initial independent set is generated from a moderate-size subset using graph-aware search. Second, additional sequence pairs are collected if they are compatible with this initial independent set. Third, a final graph-aware search on the collected sequence pairs identifies additional compatible sequence pairs, which are added to the initial independent set.

Some criteria are intrinsic properties of individual sequence pairs and can therefore be evaluated before considering interactions with other sequence pairs. These criteria include the on-target interaction between a strand and its intended binding partner, as well as the self-folding free energies of the individual strands. Applying these criteria yields a prefiltered pool of sequence pairs that are eligible for inclusion in an orthogonal sequence library. Details of the sequence sampling and prefiltering implementation are provided in Sections 6.2 and 6.3 of the Supporting Information.

Once this prefiltered pool has been defined, compatibility between sequence pairs is determined by their off-target interactions. For each comparison, four off-target interactions are possible: each strand from one sequence pair can interact with each strand from the other sequence pair. If any of these interactions exceeds the chosen off-target limit, the two sequence pairs are considered incompatible. This produces a list of all combinations of sequence pairs, with each combination classified as either compatible or incompatible. This classification can be represented as a conflict graph, where vertices represent sequence pairs and edges indicate that two sequence pairs are incompatible. In this graph, an orthogonal set of sequence pairs corresponds to a subset of vertices with no conflict edges between them. Such a subset is called an independent set. The selection problem can therefore be interpreted as finding the largest possible independent set in the conflict graph, a well-known NP-hard graph problem (*24*).

### Graph-aware search algorithm

A graph-aware heuristic strategy for finding large, but not necessarily maximum-size, independent sets is illustrated in Figure 2B. In this strategy, each vertex in the conflict graph is labeled by its number of edges, corresponding to its number of conflicts. The vertex with the highest number of edges is removed first. The number of edges is then recounted for the remaining vertices, and the procedure is repeated. Once no edges remain, the remaining vertices form an independent set. This basic removal strategy is known as the GMAX maximum-degree deletion heuristic (*20*).

Graph heuristics often improve simple greedy procedures through tie-breaking rules, local search, restart strategies, or adaptive refinement steps (*25* –*28*). Following this general strategy, we supplemented the basic removal procedure with an overlap-based tie-breaking rule, a cleanup step that reintroduces compatible excluded sequence pairs, and an iterative perturbation step that explores nearby independent sets. Details of the graph-aware search implementation are provided in Section 6.1 of the Supporting Information.

The graph-aware heuristic described above requires the complete conflict graph to be known before the search begins. This becomes impractical for longer core binding domains because the number of possible cores scales as *N* ∝ 4*^L^*, and constructing the graph requires testing all *N* (*N* − 1)*/*2 combinations of sequence pairs. Since each comparison involves four possible off-target interactions and each interaction requires a computationally expensive NUPACK calculation, full graph construction from the complete sequence space becomes impractical for core binding domains longer than 7 nucleotides.

### Sequential candidate-addition algorithm (naive search)

A commonly used alternative that avoids constructing the full conflict graph is to build the independent set sequentially (*16*), as illustrated in Figure 2C. Starting from the prefiltered pool, one random sequence pair is first added to initialize the independent set. The remaining sequence pairs are then considered one by one. A sequence pair is added only if it has no off-target conflict with any sequence pair already in the independent set. This strategy scales more favorably for large sequence-pair pools because it tests only interactions with the independent set rather than all combinations of sequence pairs. We refer to this strategy as naive search because it does not use information about the global conflict structure. Implementation details are provided in Section 6.4 of the Supporting Information.

### Hybrid search algorithm

In Figure 2D, we present a hybrid strategy that combines graph-aware elements with the favorable scaling of naive search. First, an initial subset of moderate size is selected from the prefiltered pool. Graph-aware search is applied to this subset to obtain an initial independent set. Additional sequence pairs are then considered one by one and tested against this initial independent set. Sequence pairs with no off-target conflict with the initial independent set are added to a collection pool. Sequence pairs in the collection pool are therefore compatible with the initial independent set, but not necessarily with each other. After the collection stage, a final graph-aware search is performed on the collection pool to identify an additional independent set. Because these additional sequence pairs are compatible with the initial independent set by construction, they can be added to it to obtain a larger independent set. The size of the subset used for the initial graph-aware search is a user-defined parameter. Accordingly, graph-aware search can be implemented as a special case of the hybrid workflow by setting the initial subset to the desired size and omitting the subsequent collection and final graph-aware search stages. In this configuration, the independent set obtained from the initial graph is returned directly as the final sequence-pair library. Implementation details are provided in Section 6.5 of the Supporting Information.

### Graphical user interface (OrthoSeq)

Creating an orthogonal sequence-pair pool requires several choices that depend on the intended experimental conditions, including temperature, salt concentration, core binding-domain length, optional 5*^′^*and 3*^′^* extensions, and the energy limits used for sequence selection, as illustrated in Figure 1. Because these choices are interdependent and difficult to adjust manually, we implemented the graphical user interface illustrated in Figure 3. The interface guides the user through sequence-layout definition (Figure 3A), selection of the on-target energy range (Figure 3B), self-folding energy limit (Figure 3C), and off-target energy limit (Figure 3D), and generation of the final orthogonal library using hybrid search (Figure 3E).

**Figure 3:**
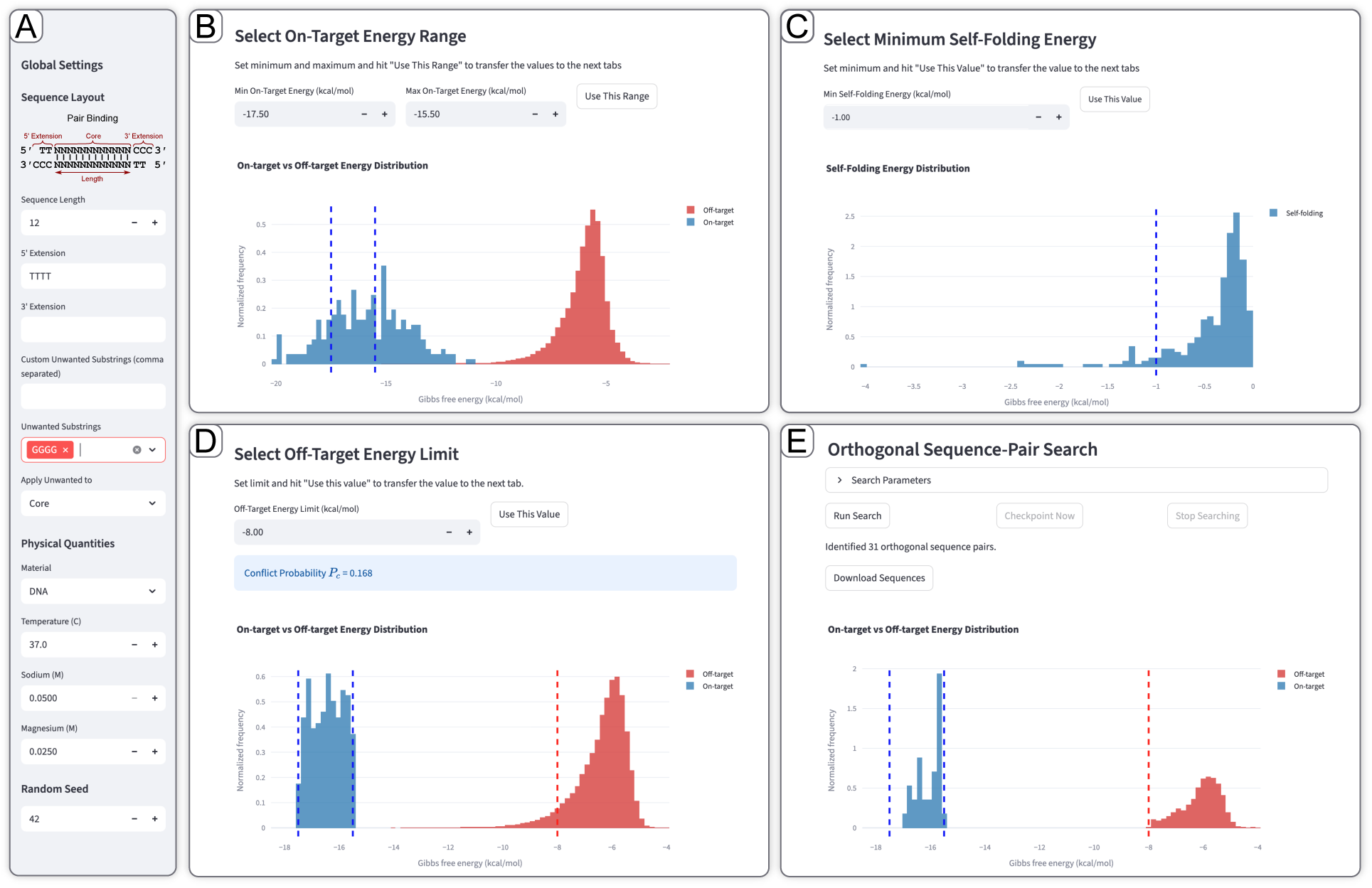
Graphical user interface for custom sequence-pair library generation. **(A)** Sequence-layout and physical-parameter selection. The user defines the core binding-domain length, optional 5*^′^* and 3*^′^*extensions, excluded motifs, and NUPACK parameters. **(B)** On-target and off-target association free-energy distributions used to select the on-target energy range. User-selected limits entered in the input fields are displayed as vertical lines. **(C)** Self-folding free energy distribution used to select the self-folding energy limit. The user-selected limit entered in the input field is displayed as a vertical line. **(D)** Refined on-target and off-target association free-energy distributions after applying the selected on-target and self-folding energy limits. The user-selected off-target energy limit is displayed as a vertical line and used to calculate the conflict probability, which is shown in a blue box. **(E)** Search panel used to configure and run the hybrid search algorithm. After completion, the final energy distributions are displayed and the resulting sequence-pair library can be saved as an Excel report.

The workflow begins with the definition of the sequence layout and the physical parameters used for the NUPACK calculations. These include the core binding-domain length, optional 5*^′^*and 3*^′^* extensions, excluded sequence motifs, nucleic acid type, temperature, sodium concentration, and magnesium concentration (Figure 3A).

A subset of sequence pairs satisfying the defined sequence layout is then sampled. Their on-target and off-target association free energies are calculated and plotted as distributions, allowing the user to select an appropriate on-target energy range (Figure 3B). The self-folding free energy distribution is displayed separately to allow selection of the self-folding energy limit (Figure 3C). A selection-helper view, not shown in Figure 3, relates the association and self-folding free energies to the predicted bound and unpaired fractions described above, allowing the limits to be chosen using physically interpretable criteria.

In the next step, new sequence pairs that satisfy the selected on-target energy range and self-folding energy limit are sampled, and their on-target and off-target association free energies are calculated. The resulting energy distributions are displayed so that the final off-target energy limit can be chosen (Figure 3D). This step is separated from the initial on-target selection because the on-target and off-target distributions are not independent. Sequence pairs with stronger on-target binding generally also show stronger off-target binding, so changing the allowed on-target range changes the off-target distribution of the remaining sequence-pair pool.

The selected parameters are then used to run the hybrid search algorithm (Figure 3E). Search-specific parameters, including the initial graph-aware search subset size, the number of graph-aware search iterations, and the perturbation fraction, can be adjusted before starting the search. After completion, the resulting orthogonal sequence-pair library can be exported together with the corresponding search parameters. The interface also allows users to initialize the workflow from seqwalk-generated candidate pools instead of generating candidate sequences *de novo*. OrthoSeq was implemented and tested in a Conda environment using Python 3.11.14 and NUPACK 4.0.2.0. Additional implementation details are provided in Section 6.7 of the Supporting Information.

### Benchmark design

To compare the search strategies, we benchmarked the size of the orthogonal sequence-pair sets obtained under fixed thermodynamic and sequence design constraints for several core binding-domain lengths, with and without a 5*^′^* TTTT extension. The extension represents a commonly used flexible linker and allowed us to assess how such a linker affects the thermodynamic constraints and sequence-search performance. We separated the benchmarks into two regimes. For short core binding domains, full conflict-graph construction is computationally feasible, allowing direct comparison of graph-aware search, naive search, and hybrid search. For longer core binding domains, full graph construction becomes impractical, so the strategies were compared under a fixed computational budget of 10 million NUPACK calls, corresponding to approximately 20 h of CPU computation on a MacBook Air with an Apple M1 chip, 8 GB of memory, and macOS 14.8.7. For hybrid search, we additionally varied the size of the random subset used for the initial graph-aware search and tested subsets of 250, 450, and 900 sequence pairs. We also included a graph-aware search condition in which the full computational budget was allocated to the initial graph construction and search, resulting in an initial graph containing approximately 2235 sequence pairs.

Because binding strength depends strongly on core binding-domain length, the allowed on-target energy range was chosen separately for each length from the corresponding on-target energy distribution. Specifically, the range was defined from the mean on-target energy to one standard deviation below the mean. The off-target limit was chosen differently for short and long core binding domains. For longer core binding domains, we used a bound-fraction limit requiring the predicted off-target bound fraction to remain below 1 % at 1 *µ*M strand concentration, corresponding to −8.16 kcal*/*mol.

For short core binding domains, this limit is less informative because on-target binding is already weak and most off-target interactions satisfy the same bound-fraction limit automatically. We therefore adjusted the off-target limit separately for each condition. The off-target limit was chosen to produce conflict probabilities of 0.1, 0.2, and 0.3 in the prefiltered input pool before graph-aware or hybrid search was applied. The conflict probability was defined as the probability that two randomly chosen sequence pairs from this pool were incompatible.

For all benchmarks, the self-folding energy limit was chosen such that the predicted unpaired fraction remained above 0.2, corresponding to −0.99 kcal*/*mol. Other physical parameters and sequence design constraints were set as shown in Figure 1B. Further benchmark implementation details are provided in Section 6.6 of the Supporting Information. The short sequence off-target limits are listed in Supplementary Table S1.

### Seqwalk comparison datasets

To place the thermodynamic-search results in context, we compared them with sequence libraries generated by seqwalk. For the seqwalk-only comparison in Figure 5A, we generated 16-nucleotide barcodes with seqwalk using *k* = 6, where *k* denotes the length of the sequence words used by the seqwalk exclusion rule. We used the maximum-orthogonality setting, a requested library size of at least 50 sequences, and a GC-content constraint of 7 to 11 G or C bases per sequence. This yielded 72 barcode sequences. Each barcode was paired with its reverse complement to define a sequence pair. No 5*^′^* or 3*^′^* extensions were used. Sequences containing GGGG or CCCC motifs were excluded to avoid G-quadruplex-forming sequences. The resulting sequence pairs were evaluated with NUPACK to calculate on-target association energies, off-target association energies, and self-folding energies.

For the direct comparison in Figure 5B, we used the 16-mer sequence-pair library obtained from the long-sequence benchmark shown in Figure 4. This library was generated with the hybrid search algorithm using an initial subset size of 450 and the benchmark thermodynamic limits described above. The final library contained 74 sequence pairs.

**Figure 4:**
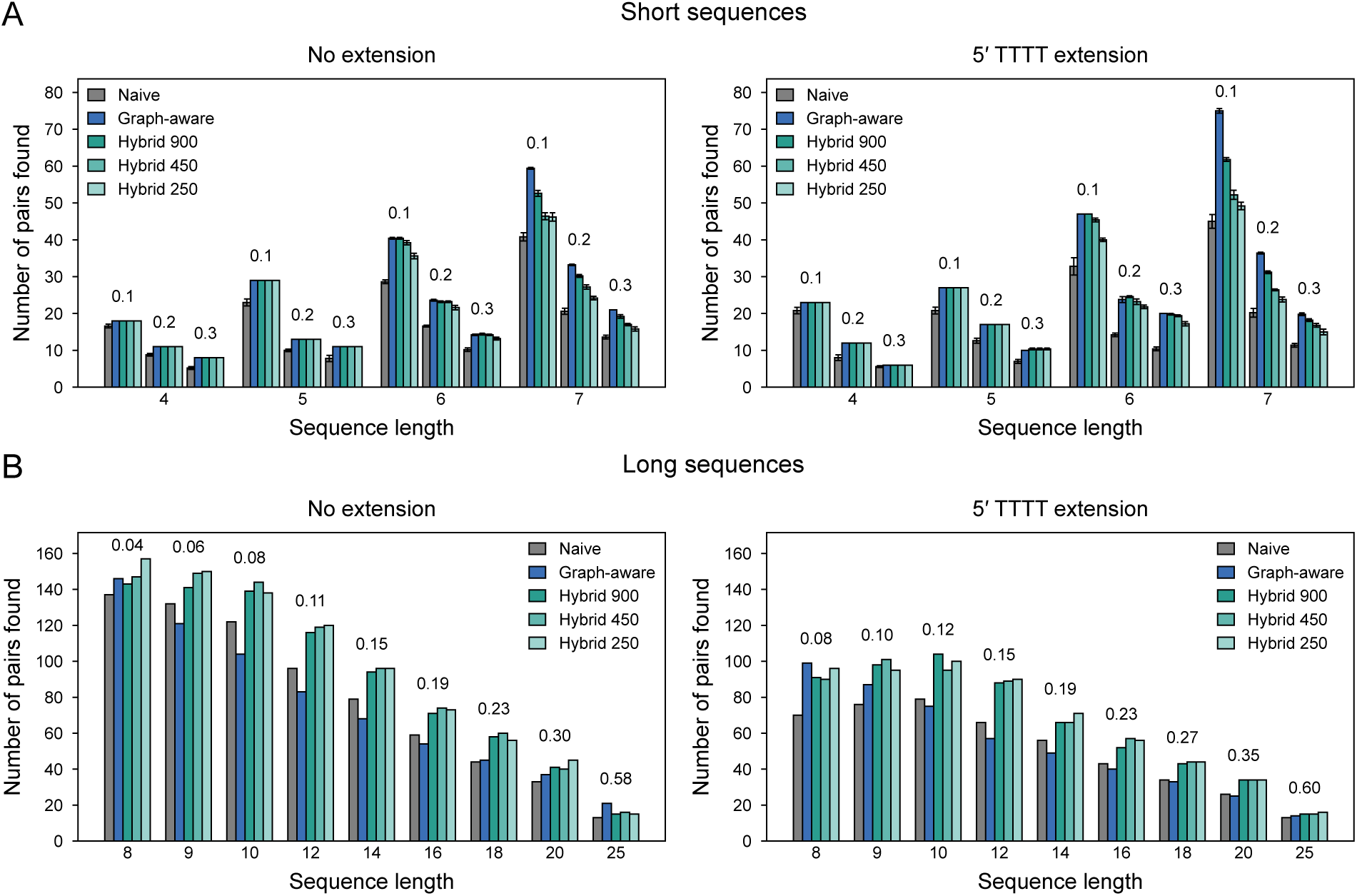
Benchmark comparison of sequence-pair search strategies. **(A)** Short sequence benchmark. Number of orthogonal sequence pairs found by naive search, hybrid search, and graph-aware search for short core binding domains, with and without a 5*^′^* TTTT extension. For hybrid search, the numbers in the legend indicate the subset size used for the initial graph-aware search. Benchmarks were performed at conflict probabilities of 0.1, 0.2, and 0.3, which are indicated above the corresponding bar clusters. Each bar represents five runs with different random seeds, and error bars show the standard deviation. **(B)** Long sequence benchmark. Number of orthogonal sequence pairs found by naive search, hybrid search, and graph-aware search under a fixed computational budget of 10 million NUPACK calls. For hybrid search, the numbers in the legend indicate the subset size used for the initial graph-aware search. For the graph-aware search condition, the initial subset size was chosen to use the full computational budget, corresponding to approximately 2235 sequence pairs. Conflict probabilities for each condition are indicated above the bars.

**Figure 5:**
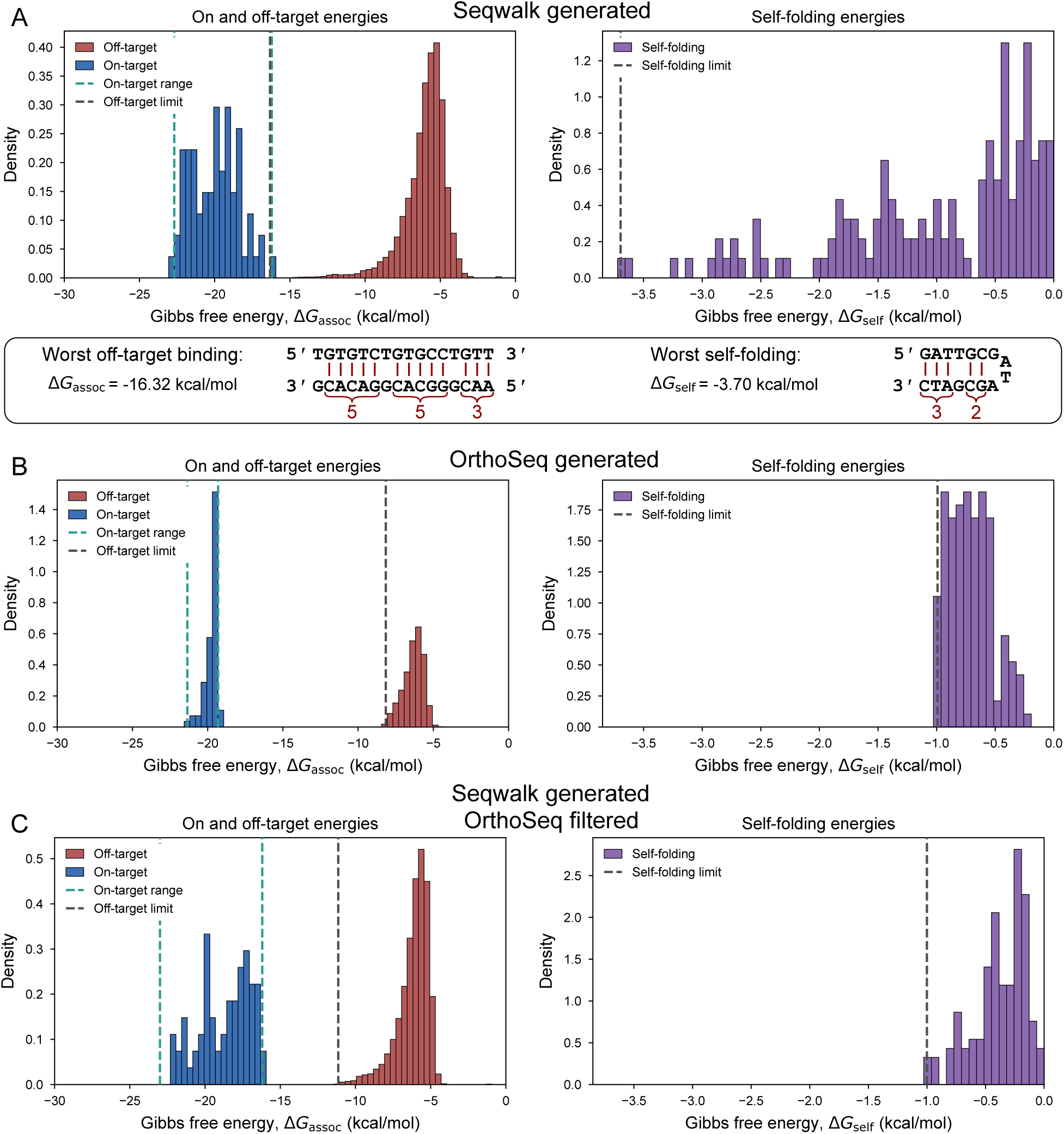
Thermodynamic comparison of seqwalk-generated and thermodynamically selected sequence libraries. **(A)** Energy distributions for 72 sequence pairs generated from 16-nucleotide seqwalk barcodes using *k* = 6, a GC-content constraint of 7 to 11 G or C bases, and the maximum-orthogonality setting. The strongest off-target interaction and the sequence with the strongest self-folding are illustrated below the corresponding distributions. **(B)** Energy distributions for the 74 sequence pairs from the 16-mer long-sequence benchmark with an initial subset size of 450, shown in Figure 4. **(C)** Energy distributions for 72 sequence pairs obtained after thermodynamic postfiltering of a seqwalk-generated candidate pool.

For the seqwalk postfiltering comparison in Figure 5C, we generated a predefined candidate pool with seqwalk using barcode length *L* = 16 and *k* = 6, where *k* again denotes the length of the sequence words used by seqwalk. Reverse-complement avoidance and GC-content constraints were not imposed. Sequences containing GGGG or CCCC motifs were excluded during generation by seqwalk. Each seqwalk barcode was paired with its reverse complement to define a sequence pair. The candidate pool was then filtered using the same thermodynamic criteria used throughout this work. The allowed on-target association-energy range was set to −23.0 to −16.2 kcal*/*mol, the off-target limit was set to −11.143 kcal*/*mol, and the self-folding energy limit was set to −1.0 kcal*/*mol. The initial subset size was set to 900, with a perturbation fraction of 0.2 and 1000 graph-aware search iterations. Because the seqwalk-derived candidate pool was smaller than the requested initial subset, only the initial graph-aware search step was performed. The final postfiltered library contained 72 sequence pairs.

## Results

### Search algorithm benchmarks

We first evaluated whether graph-aware search and hybrid search find larger orthogonal sequence-pair libraries than sequential candidate addition, referred to here as naive search, under the benchmark conditions defined above. The results of the short sequence (4–7 bases per core binding domain) benchmark are shown in Figure 4A. Graph-aware search and hybrid search outperformed naive search under all tested conditions. The improvement was largest for 6-mer and 7-mer core binding domains and was more pronounced for sequence pairs containing the 5*^′^*TTTT extension. The strongest improvement was observed for 6-mer core binding domains with a 5*^′^*TTTT extension at a conflict probability of 0.3, where graph-aware search found almost twice as many sequence pairs as naive search. As expected, graph-aware search produced the largest sequence-pair libraries under all tested conditions, while hybrid search either matched or slightly trailed its performance. For hybrid search, larger initial subset sizes generally increased the number of sequence pairs found because the method becomes more similar to full graph-aware search as the initial subset size increases. For 4-mer and 5-mer core binding domains, graph-aware and hybrid search gave identical results because the prefiltered pools contained only 37 and 163 sequence pairs, respectively, both smaller than the tested initial subset sizes.

The long sequence benchmark for core binding domains longer than 7 nucleotides is shown in Figure 4B. Hybrid search with initial subset sizes of 250, 450, and 900 outperformed naive search for all tested core binding-domain lengths, both with and without the 5*^′^* TTTT extension. In contrast, the graph-aware search condition, in which the entire computational budget was used to construct the initial graph, generally performed worse than naive search or approximately the same. Exceptions were observed for the shortest sequences with core binding-domain lengths of 8 and 9 nucleotides and for the longest sequences with a core binding-domain length of 25 nucleotides and no 5*^′^*extension.

For the 8-nucleotide condition without an extension and the 8- and 9-nucleotide conditions with the 5*^′^* TTTT extension, graph-aware search outperformed naive search. For the 8-nucleotide condition with the 5*^′^* TTTT extension, it also outperformed the tested hybrid-search conditions. This behavior can be explained by the fact that, in these cases, naive search and hybrid search with initial subset sizes of 250, 450, and 900 exhausted the available sequence pool before reaching the computational limit, whereas the graph-aware search condition used the entire computational budget to construct and search the initial graph.

The difference between naive and graph-aware search progressively decreased as the core binding-domain length increased, and the two approaches produced comparable results from 18 nucleotides onward. At 25 nucleotides without a 5*^′^* extension, graph-aware search outperformed naive search and all tested hybrid-search conditions. However, this crossover was not observed for the corresponding condition with the 5*^′^* TTTT extension. For intermediate core binding-domain lengths, graph-aware search generally produced smaller libraries than naive search, whereas the hybrid-search conditions still produced meaningfully larger libraries than both. This shows that allocating the full computational budget to graph construction on a restricted subset was generally less effective than combining a smaller graph-aware initialization with exploration of additional candidates, although the relative performance of graph-aware search improved at longer core lengths.

The relative improvements achieved by hybrid search over naive search in the long-sequence benchmark were smaller than those achieved by graph-aware search over naive search in the short-sequence benchmark. Without a 5*^′^* extension, hybrid search found approximately 15–25 % more sequence pairs than naive search. With the 5*^′^* TTTT extension, the improvement was approximately 20–30 %. Among the hybrid-search conditions with initial subset sizes of 250, 450, and 900, no clear dependence on initial subset size was observed. Conflict probability increased with core binding-domain length, consistent with the larger number of possible off-target interactions for longer sequences. Correspondingly, the number of sequence pairs found generally decreased with increasing conflict probability. For the same core binding-domain length, sequence pairs with the 5*^′^* TTTT extension had higher conflict probabilities and yielded smaller orthogonal sets, indicating that the extension introduced additional off-target interactions. We repeated the long-sequence benchmark at 25 *^◦^*C without the graph-aware search condition and obtained similar results for naive and hybrid search (Supplementary Figure S3).

Overall, hybrid search with an initial subset size of 450 appears to provide the best practical choice for longer sequences. It consistently produced competitive results and showed no meaningful disadvantage compared with the other tested hybrid-search conditions, although no clear correlation between initial subset size and final library size was observed. For short sequences with core binding domains up to 7 nucleotides, full graph-aware search appears to be the optimal strategy. For each benchmark condition, the largest sequence-pair library generated across the replicate runs is available through Zenodo as a ready-to-use sequence set (*29*).

To gain further insight into the benchmark results, we analyzed the conflict structure of representative 12-mer sequence-pair graphs in Section 3 of the Supporting Information. The conflict probability varied substantially between sequence pairs. This indicates that the conflict graph is not well described by a simple random-graph model in which every edge occurs independently with the same probability (commonly known as an Erdös-Rényi graph (*30*)). This variation suggests that some sequence pairs are intrinsically more conflict-prone than others. Sequence pairs obtained by naive search had fewer conflicts with additional sampled candidate sequence pairs than randomly selected sequence pairs. Sequence pairs obtained by the initial graph-aware search of the hybrid algorithm had even fewer conflicts, indicating stronger enrichment in globally low-conflict sequence pairs. Together, these results suggest that both algorithms enrich the selected independent set in globally low-conflict sequence pairs, even though each algorithm evaluates only part of the full conflict graph. Both strategies also showed approximately logarithmic growth in the number of sequence pairs found as the number of considered sequence pairs increased.

### Comparison with seqwalk-generated sequence libraries

We also compared our thermodynamic sequence-selection workflow with seqwalk, a recently introduced method for generating large DNA barcode libraries using sequence-symmetry minimization (*19*). This comparison allowed us to determine how sequence-level heuristics perform under thermodynamic evaluation and whether the resulting libraries can be improved by thermodynamic postfiltering with OrthoSeq. Seqwalk reduces similarity between barcodes by preventing short sequence words of length *k* from appearing more than once in the library. In the maximum-orthogonality setting, the reverse complements of these sequence words are also excluded. Seqwalk additionally allows the GC content of generated sequences to be restricted, which can make binding strength more predictable. These criteria are useful for barcode design, but they are not equivalent to thermodynamic criteria because binding interactions are not evaluated directly.

We first evaluated a seqwalk maximum-orthogonality library using the same NUPACK-based thermodynamic criteria used throughout this work. The resulting energy distributions are shown in Figure 5A. Although the sequences satisfy the seqwalk sequence-level orthogonality criterion by prohibiting repeated and reverse-complementary 6-nucleotide words, they are not thermodynamically orthogonal. The strongest off-target interaction had Δ*G*_assoc_ = −16.32 kcal*/*mol, which was slightly stronger than the weakest on-target interaction, Δ*G*_assoc_ = −16.2 kcal*/*mol. Several sequences also showed unfavorable self-folding.

The strongest off-target interaction and the sequence with the strongest self-folding are illustrated below the energy distributions in Figure 5A. The strongest off-target interaction contains two complementary 5-bp stretches and one additional 3-bp complementary stretch. Seqwalk suppresses complementary 6-mers in the maximum-orthogonality setting, but it does not exclude multiple shorter complementary regions separated by gaps. Similarly, the sequence with the strongest self-folding contains a hairpin stabilized by separated 3-bp and 2-bp complementary regions. These examples show why *k*-mer orthogonality and thermodynamic orthogonality are related but not equivalent.

For comparison, Figure 5B shows the energy distributions for the 16-mer sequence-pair library obtained from the long-sequence benchmark in Figure 4. In contrast to the seqwalk-only library, the on-target energies occupy a narrow range, all off-target interactions remain above the imposed limit, and the self-folding energies remain above the selected limit. Thus, this library provides a direct comparison for the thermodynamic quality obtained with our search workflow.

Finally, we tested whether seqwalk-generated candidates could be used as an input pool for thermodynamic refinement. A seqwalk-derived candidate pool was filtered with the same thermodynamic sequence-selection workflow, using the limits described in Methods. This postfiltering strategy produced 72 thermodynamically orthogonal sequence pairs, shown in Figure 5C.

Together, these comparisons show that seqwalk alone can generate sequence-level barcode libraries that still contain substantial thermodynamic cross-hybridization. Our workflow can generate thermodynamically constrained sequence-pair libraries directly, or it can refine predefined seqwalk candidate pools when preserving barcode-like sequence properties is desired.

## Discussion

### Isolated binding reactions as conservative design criteria

The isolated two-strand description used to define the free-energy limits does not represent the full experimental pool, where all sequences are present simultaneously and compete through many possible interactions. However, this simplification provides a conservative criterion for off-target binding. If a specific off-target interaction is weak in isolation, the same interaction will be at least as weak in the full pool, because additional competing interactions can only reduce the fraction of material bound in that specific off-target complex.

### Thermodynamic limits and library size

The thermodynamic limits used in the benchmarks were kept fixed to allow direct comparison across sequence lengths and search strategies. Under these conditions, the number of sequence pairs found decreased with increasing core binding-domain length because longer sequences produced more off-target conflicts. Larger libraries could be obtained by using a less stringent off-target energy limit. For longer sequences, this may be reasonable because the energetic gap between on-target and off-target binding generally increases with sequence length. Whether this is acceptable depends on the application. If on-target binding only needs to outcompete off-target binding, a weaker off-target restriction may be sufficient. If transient off-target binding must also be avoided, the off-target limit must remain stringent. Larger libraries could also be obtained by changing the experimental conditions, for example by increasing the temperature or relaxing the self-folding criterion.

### Conflict structure and search algorithms

The conflict graphs are heterogeneous, with some sequence pairs being intrinsically more conflict-prone than others (Supporting Information, Section 3). Both naive search and graph-aware search select sequence pairs that are globally less conflict-prone than randomly sampled candidate sequence pairs, with graph-aware search doing so more effectively. This helps explain the advantage of hybrid search. In naive search, conflict-prone sequence pairs can enter the independent set early and then reduce the probability of accepting additional sequence pairs in later steps. In hybrid search, the initial independent set is generated by graph-aware search and is therefore enriched in globally low-conflict sequence pairs. This makes it easier to identify additional sequence pairs during the subsequent collection stage. At the same time, the competitive performance of naive search for longer core binding domains under a fixed computational budget can be explained by the fact that sequential rejection also enriches the independent set in globally low-conflict sequence pairs, although less effectively than graph-aware search.

### Computational scaling

For short core binding domains, full conflict-graph construction is feasible and allows the search to use the complete conflict structure. For longer core binding domains, however, the number of possible sequence pairs grows rapidly with core length, and constructing a conflict graph for *T* candidate sequence pairs requires testing *T* (*T* − 1)*/*2 combinations. Graph construction therefore becomes the limiting computational cost.

Both sequential candidate addition and full conflict-graph search showed approximately logarithmic scaling in the number of sequence pairs found with the number of considered candidate sequence pairs (Supporting Information, Section 3). To interpret this behavior, we estimated the scaling of both strategies for a simplified conflict graph with uniform conflict probability in Section 2 of the Supporting Information. We found that the independent-set size scales as *I*(*T*) ∼ *α* ln *T* at leading order, where *T* is the number of considered candidate sequence pairs. In this estimate, the prefactor *α* is at most twofold larger for full conflict-graph search than for sequential candidate addition.

However, this advantage is offset by the computational cost of graph construction. Denoting the computational cost by *C*, naive search scales approximately linearly with the number of considered candidates *T*, *C*_naive_ ∼ *T*, whereas full graph construction scales quadratically, *C*_graph_ ∼ *T* ^2^. Consequently, the up to twofold larger logarithmic prefactor estimated for graph-aware search is at best canceled when the scaling is expressed as a function of computational cost. Thus, naive search scales more favorably with computational cost than full graph-aware search when conflict-edge construction is limiting and edge probabilities are assumed to be uniform.

The benchmark results are mostly consistent with this estimate. For intermediate core binding-domain lengths, graph-aware search generally produced smaller libraries than naive search. However, the difference progressively decreased as the core binding-domain length increased, and the two approaches produced comparable results from 18 nucleotides onward. We suspect that this trend arises because increasing core length produces more conflicts, making the global conflict structure increasingly important for sequence-pair selection and thus favoring graph-aware search, which exploits this structure more effectively. At 25 nucleotides without a 5*^′^* extension, graph-aware search outperformed naive search and all tested hybrid-search conditions. Although this continues the overall trend toward improved graph-aware performance at longer lengths, the absence of the same effect with the TTTT extension suggests additional condition-specific features of the conflict graph.

### Rationale behind hybrid search

Hybrid search was designed around the insights from the computational-scaling estimates and the heterogeneous conflict structure of the graph. The initial graph-aware search exploits this conflict structure by biasing the initial independent set toward globally low-conflict sequence pairs. The collection stage then uses most of the computational budget to exploit the favorable scaling of sequential candidate testing against this fixed initial independent set. Finally, graph-aware search is applied again to recover as many mutually compatible sequence pairs as possible from the sequence pairs obtained during the collection stage.

Hybrid search is also well suited for parallel computing. In the current implementation, the initial graph-search subset can be generated and evaluated as a fixed batch of sequence-pair comparisons. The collection stage could also be parallelized because the initial independent set remains fixed during this stage and can be passed to independent workers. In contrast, naive search is harder to parallelize efficiently because each newly accepted sequence pair changes the growing independent set against which later candidates must be tested.

### Thermodynamic post-selection of libraries with sequence-level orthogonality

The seqwalk comparison illustrates how OrthoSeq can complement efficient sequence-level methods for library generation. Seqwalk generated a large library satisfying the intended sequence-symmetry constraints, while thermodynamic evaluation identified off-target interactions comparable in strength to intended on-target interactions and sequences with unfavorable self-folding. OrthoSeq can therefore be used to post-select seqwalk-generated or otherwise predefined candidate pools using NUPACK-based thermodynamic criteria when predictable hybridization behavior or additional constraints are required.

Notably, a related limitation was observed in Sidewinder assembly (*13*), where auxiliary sequence pairs guide fragment ligation through DNA three-way junctions. Sidewinder used either pre-generated orthogonal barcode sets from DropSynth or custom auxiliary sequences selected with NUPACK for the intended three-way-junction context (*31*). The DropSynth barcodes were generated by removing self-dimer-forming candidates and applying sequence-level constraints on GC content, melting temperature, and modified Levenshtein distance (*31*). Assemblies using the pre-generated barcode sets showed higher misligation than assemblies using custom NUPACK-designed auxiliary sequences, further illustrating that sequence-level orthogonality does not necessarily guarantee thermodynamic orthogonality in a specific experimental context (*13*).

### Concluding remarks

We developed OrthoSeq, a workflow for generating DNA sequence-pair libraries with thermodynamically defined orthogonality. Intended binding, off-target binding, and self-folding are evaluated with NUPACK under user-defined experimental conditions, allowing binding behavior to be specified directly rather than through sequence-level heuristics alone.

Sequence pairs that violate the off-target criterion define conflicts, which allows sequence selection to be interpreted as an independent-set problem in a conflict graph. In this setting, the computational bottleneck is generating the graph, because each edge requires thermodynamic evaluation of off-target binding. Our hybrid search algorithm addresses this regime by combining an initial low-conflict set with computationally efficient sequential candidate addition, and consistently found larger sequence-pair libraries than the commonly employed sequential candidate-addition strategy.

The graphical user interface makes OrthoSeq broadly accessible for custom library generation. The seqwalk comparison shows that sequence-level barcode orthogonality does not necessarily imply thermodynamic orthogonality, while seqwalk-generated candidate pools can still be refined when predictable hybridization behavior is required. More generally, the underlying software can be used either for *de novo* sequence-pair generation or for thermodynamic refinement of predefined candidate libraries generated by other sequence design tools or the requirements of a particular experimental system. This makes OrthoSeq useful for applications requiring many mutually compatible DNA binding pairs, including DNA nanostructure assembly (*1* –*3*), multiplexed imaging (*9*, *10*), DNA-PAINT (*11*, *12*), strand-displacement systems (*7*), and other programmable hybridization-based technologies.

## Data Availability

The OrthoSeq sequence-selection workflow and the analysis scripts used in this study are available on GitHub as part of the *crisscross-kit* Python library (*32*). The benchmark data are archived on Zenodo (*29*), together with convenience copies of the largest sequence-pair library obtained for each benchmark condition in a directly usable format. OrthoSeq relies on NUPACK for thermodynamic calculations. NUPACK must be obtained and installed separately according to its licensing and installation requirements.

## Acknowledgements

The O2 High Performance Compute Cluster, supported by the Research Computing Group at Harvard Medical School, was highly beneficial in generating the data for this work. In particular, the O2 cluster was used to accelerate development of the search algorithms and to perform the final large-scale parameter sweeps. OpenAI Codex and Anthropic Claude Code were used to assist with software development. ChatGPT and Claude were used for language editing, sentence phrasing, and typographical corrections. All generated code and text were reviewed and validated by the authors.

## Author Contributions

F.K. conceived the study, developed the algorithms and software, performed the computational analyses, prepared the figures, acquired funding, and wrote the original manuscript. M.A. contributed to software development and computational analyses, acquired funding, and reviewed and edited the manuscript. W.M.S. supervised the project, acquired funding, and reviewed and edited the manuscript. All authors approved the final manuscript.

