## Supporting Information for "OrthoSeq: A Design Workflow for Thermodynamically Orthogonal DNA Sequence-Pair Libraries"

**4 Perturbation fraction analysis** **16**

**5 Long sequence benchmark at 25 °C** **17**

**6 Supplementary Methods** **19**

**7 Supplementary Tables** **26**

**References** **27**

### 1 Energy-threshold calculations

#### 1.1 Heterodimer formation

We quantify the binding strength of two strands $A$ and $B$ by considering the association reaction $A + B \rightleftharpoons AB$. The fraction of strand $A$ bound in the complex is defined as $P_b = [AB]/[A]_0$, where $[A]_0$ is the total concentration of strand $A$. For this reaction, the concentration-based equilibrium constant is

$$K = \frac{[AB]}{[A][B]}, \tag{1}$$

where $[A]$ and $[B]$ are the unbound strand concentrations at equilibrium.

We define the Gibbs free energy of association in accordance with the NUPACK thermodynamic model (*1–4*) as

$$\Delta G_{\text{assoc}} = G_{AB} - G_{\text{self},A} - G_{\text{self},B}. \tag{2}$$

Here, $G_{AB}$ is the ensemble free energy of the two-strand complex, and $G_{\text{self},A}$ and $G_{\text{self},B}$ are the single-strand ensemble free energies of strands $A$ and $B$, respectively.

The equilibrium constant is related to $\Delta G_{\text{assoc}}$ by

$$K = \frac{\exp\left(-\Delta G_{\text{assoc}}/RT\right)}{\rho_{\mathrm{H_2O}}}, \tag{3}$$

where $R$ is the gas constant, $T$ is the absolute temperature, and $\rho_{\mathrm{H_2O}} = 55.14\,\mathrm{mol\,L^{-1}}$ is the molarity of water. The factor $\rho_{\mathrm{H_2O}}$ converts the mole-fraction convention used by NUPACK to molar concentrations.

To compute thresholds, we assume equal total strand concentrations, $[A]_0 = [B]_0$. At equilibrium, the unbound strand concentrations are $[A] = [A]_0 - [AB]$ and $[B] = [A]_0 - [AB]$. Substituting these concentrations into the definition of $K$ gives

$$K = \frac{[AB]}{\left([A]_0 - [AB]\right)^2}. \tag{4}$$

Using $P_b = [AB]/[A]_0$, this can be written as

$$K[A]_0 = \frac{P_b}{(1 - P_b)^2}. \tag{5}$$

Solving for $P_b$ gives

$$P_b = \frac{(2\alpha + 1) - \sqrt{1 + 4\alpha}}{2\alpha}, \qquad \alpha = K[A]_0. \tag{6}$$

Thus, for a given strand concentration $[A]_0$, temperature $T$, and association free energy

$\Delta G_{\text{assoc}}$, the expected bound fraction can be computed directly from the equilibrium model above. Conversely, this relationship can be used to choose an association-free-energy threshold corresponding to a desired bound-fraction limit.

#### 1.2 Self-folding

We also quantify the tendency of each individual strand to form intramolecular secondary structures using the single-strand ensemble free energy $G_{\text{self}}$. This value is computed from the NUPACK partition function over all self-folding configurations of an individual strand. More negative values of $G_{\text{self}}$ correspond to a stronger tendency to self-fold.

Conceptually, self-folding can be viewed as a transition between the fully unpaired strand $U$ and the ensemble of all other folded configurations $S$,

$$U \rightleftharpoons S. \tag{7}$$

NUPACK uses the fully unpaired state as the reference state with $G_0 = 0$. The full single-strand partition function includes both the fully unpaired state and all folded states and is related to the ensemble free energy by

$$Q = \exp\left(-G_{\text{self}}/RT\right). \tag{8}$$

The Boltzmann weight of the fully unpaired reference state is $Q_0 = \exp\left(-G_0/RT\right) = 1$. The probability of the fully unpaired state is therefore

$$P_u = \frac{Q_0}{Q} = \frac{\exp\left(-G_0/RT\right)}{\exp\left(-G_{\text{self}}/RT\right)}. \tag{9}$$

Since $G_0 = 0$, this simplifies to

$$P_u = \exp\left(G_{\text{self}}/RT\right). \tag{10}$$

A required minimum unpaired fraction $P_u$ can therefore be converted into a self-folding free energy threshold for $G_{\text{self}}$.

#### 1.3 Homodimer formation

The case of homodimerization, in which $A = B$, must be treated separately. In this case, a strand binds to another copy of itself according to $A + A \rightleftharpoons A_2$. The equilibrium constant is given by

$$K_{\text{homo}} = \frac{[A_2]}{[A]^2} = \frac{[A_2]}{\left([A]_0 - 2[A_2]\right)^2}. \tag{11}$$

Here, we used the modified conservation law $[A]_0 = [A] + 2[A_2]$, because each homodimer contains two copies of strand $A$. The fraction of strand $A$ bound in the homodimer is similarly defined as $P_b = 2[A_2]/[A]_0$, which gives $[A_2] = P_b[A]_0/2$. After inserting this relation into the equilibrium constant and rearranging, we obtain

$$K_{\mathrm{homo}}[A]_0 = \frac{P_b}{2\left(1 - P_b\right)^2}. \tag{12}$$

Comparing this result with equation 5 suggests defining an effective equilibrium constant $K_{\mathrm{eff}} = 2K_{\mathrm{homo}}$, which maps the homodimer relation onto the corresponding heterodimer relation. The homodimer equilibrium constant is related to the Gibbs free energy of homodimer association, $\Delta G^{\mathrm{homo}}_{\mathrm{assoc}}$, which gives

$$K_{\mathrm{eff}} = 2K_{\mathrm{homo}} = \frac{2\exp\left(-\Delta G^{\mathrm{homo}}_{\mathrm{assoc}}/RT\right)}{\rho_{\mathrm{H_2O}}} = \frac{\exp\left[-\left(\Delta G^{\mathrm{homo}}_{\mathrm{assoc}} - RT\ln 2\right)/RT\right]}{\rho_{\mathrm{H_2O}}}. \tag{13}$$

This in turn suggests defining an effective Gibbs free energy of association for homodimer formation as

$$\Delta G^{\mathrm{homo}}_{\mathrm{assoc,eff}} = \Delta G^{\mathrm{homo}}_{\mathrm{assoc}} - RT\ln 2. \tag{14}$$

Whether the uncorrected or effective homodimer association free energy should be used for thresholding depends on the physical quantity that the threshold is intended to constrain. Using $\Delta G^{\mathrm{homo}}_{\mathrm{assoc}}$ applies the same association-free-energy threshold to heterodimer and homodimer formation. In contrast, using $\Delta G^{\mathrm{homo}}_{\mathrm{assoc,eff}}$ applies the same limit to the fraction of strand material bound in either type of complex. Both interpretations can be experimentally relevant. A common association-free-energy threshold treats heterodimer and homodimer interactions with the same association strength equally, whereas a common bound-fraction threshold accounts for the fact that each homodimer removes two copies of the same strand from the free strand population. The benchmarks presented in this work were performed using the uncorrected NUPACK association free energies and therefore applied the same association-free-energy threshold to heterodimer and homodimer formation. The OrthoSeq application allows the user to choose whether homodimer energies are evaluated directly or mapped onto the common bound-fraction scale using the correction above.

### 2 Algorithmic scaling estimates

The benchmark results in Section 3 show an approximately logarithmic increase in the number of orthogonal sequence pairs with the number of pre-filtered candidate sequence pairs considered. Here, we derive simple scaling estimates to rationalize this behavior. These estimates are not intended as exact models of the sequence-design conflict graphs. Instead, we derive them by assuming the idealized case of an Erdős-Rényi conflict graph (*5*) in which conflict edges occur independently with uniform probability.

#### 2.1 Naive search

In the simplified model, each newly sampled pre-filtered candidate has an independent probability $p$ of conflicting with each sequence pair already present in the growing independent set. If the current independent set contains $I$ sequence pairs, the probability that a new candidate has no conflict with any of them is

$$P_{\text{accept}}(I) = (1-p)^I. \tag{15}$$

The probability that the next accepted sequence pair is found on the $k$-th trial is

$$P_{\text{trials}}(k \mid I) = (1 - P_{\text{accept}}(I))^{k-1}\, P_{\text{accept}}(I), \tag{16}$$

which is a geometric distribution. The expected number of trials $N(I)$ to add one more sequence pair is therefore given by

$$N(I) = \sum_{k=1}^{\infty} k\,(1 - P_{\text{accept}}(I))^{k-1}\, P_{\text{accept}}(I) = \frac{1}{P_{\text{accept}}(I)}. \tag{17}$$

Using the expression for $P_{\text{accept}}(I)$, this gives

$$N(I) = (1-p)^{-I}. \tag{18}$$

The total expected number of trials $T$ required to reach an independent set size of $I$ is therefore given by

$$T(I) = \sum_{i=0}^{I-1} N(i) = \sum_{i=0}^{I-1} (1-p)^{-i}. \tag{19}$$

This sum can be simplified using the geometric-series formula, giving

$$T(I) = \frac{(1-p)^{-I} - 1}{(1-p)^{-1} - 1}. \tag{20}$$

Solving this expression for $I$ gives

$$I = \frac{\ln\left(1 + T\frac{p}{1-p}\right)}{-\ln(1-p)}. \tag{21}$$

We can read off the limiting scaling from this expression. For increasingly large $T$, the term $1 + Tp/(1-p)$ is dominated by $Tp/(1-p)$. Therefore,

$$\ln\left(1 + T\frac{p}{1-p}\right) \approx \ln T + \ln\left(\frac{p}{1-p}\right). \tag{22}$$

The second term is constant with respect to $T$, so the large-$T$ scaling is

$$I \sim \frac{\ln T}{-\ln(1-p)}. \tag{23}$$

Thus, in this simplified Erdős-Rényi model, naive search is expected to show logarithmic growth of the independent-set size with the number of considered pre-filtered candidate sequence pairs.

#### 2.2 General independent-set scaling

For graph-aware search using the GMAX heuristic, it is harder to derive a simple scaling argument directly because the result depends on the full conflict graph and on the order in which vertices are removed. We therefore estimate the size $I$ of an independent set that can be found in an Erdős-Rényi graph with $T$ vertices and conflict probability $p$. We loosely follow the first-moment approach used for clique-size scaling in Erdős-Rényi random graphs (*5*), applied here to independent sets in a conflict graph.

An arbitrary subset of $I$ vertices has $I(I-1)/2$ possible conflict edges. For this subset to be an independent set, none of these conflict edges can be present. The probability that an arbitrary subset of $I$ vertices is independent is therefore

$$P_{\text{ind}}(I) = (1-p)^{I(I-1)/2}. \tag{24}$$

The number of possible vertex subsets of size $I$ in the full graph with $T$ vertices is $\binom{T}{I}$. Thus, the expected number of independent sets of size $I$ in the graph is

$$E(I) = \binom{T}{I}(1-p)^{I(I-1)/2}. \tag{25}$$

Independent sets of size $I$ are expected to exist as long as $E(I)$ is not much smaller than one. This gives the approximate threshold condition

$$\binom{T}{I}(1-p)^{I(I-1)/2} \approx 1. \tag{26}$$

Taking the logarithm on both sides gives

$$\ln\binom{T}{I} + \frac{I(I-1)}{2}\ln(1-p) \approx 0. \tag{27}$$

Writing the binomial coefficient explicitly gives

$$\binom{T}{I} = \frac{T!}{I!(T-I)!} = \frac{T(T-1)\cdots(T-I+1)}{I!}. \tag{28}$$

For $I \ll T$, the factors in the numerator are all approximately $T$, so

$$\binom{T}{I} \approx \frac{T^I}{I!}. \tag{29}$$

Taking the logarithm gives

$$\ln\binom{T}{I} \approx I\ln T - \ln(I!). \tag{30}$$

For $I \gg 1$, we can use Stirling's approximation for $\ln(I!)$, which gives

$$\ln\binom{T}{I} \approx I\ln T - I\ln I + I = I\left(\ln T - \ln I + 1\right). \tag{31}$$

Inserting this into equation 27 gives

$$I\left(\ln T - \ln I + 1\right) + \frac{I(I-1)}{2}\ln(1-p) \approx 0. \tag{32}$$

Since $I > 0$, we can divide by $I$ and isolate $\ln T$, giving

$$\ln T \approx \ln I - 1 - \frac{I-1}{2}\ln(1-p). \tag{33}$$

This equation cannot be solved for $I$ using standard elementary analytical expressions. However, we can read off the limiting scaling. For increasingly large $I$, the term proportional to $I-1$ outcompetes the slower-growing term $\ln I - 1$. Using $I - 1 \approx I$, this gives

$$\ln T \approx -\frac{I}{2}\ln(1-p). \tag{34}$$

Solving for $I$ gives

$$I \approx \frac{2\ln T}{-\ln(1-p)}. \tag{35}$$

Thus, the expected independent-set size in an Erdős–Rényi conflict graph grows approximately logarithmically at leading order with the number of considered pre-filtered candidate sequence pairs. Interestingly, the leading-order limiting scaling is a factor of two larger than the corresponding scaling for naive search in equation 23. This suggests that improvements over naive search are possible when an algorithm can use the global conflict structure of the graph. This comparison is only a reference estimate, because the graph considered here is a simplified Erdős-Rényi graph with independent conflicts of uniform probability, whereas the sequence-design conflict graphs are likely structured and heterogeneous.

#### 2.3 Empirical fit function

To compare the observed scaling behavior with the logarithmic estimates derived above, we use an empirical fit function for the number of obtained orthogonal sequence pairs as a function of the number of considered pre-filtered candidate sequence pairs. The naive-search estimate gives the limiting scaling $I \sim \ln T/[-\ln(1-p)]$, whereas the general independent-set estimate gives $I \sim 2\ln T/[-\ln(1-p)]$. Both estimates therefore predict logarithmic limiting scaling, but with different prefactors. We therefore require a fit function with limiting scaling $A\ln T$, where $A$ is an empirical constant. We also require the function to satisfy $I(0) = 0$, giving

$$I = A\ln\left(1 + \frac{T}{B}\right). \tag{36}$$

This function has the same mathematical form as the naive-search result in equation 21, where $A = 1/[-\ln(1-p)]$ and $B = (1-p)/p$. For the empirical fits, however, $A$ and $B$ are treated as fit parameters rather than being fixed by a single conflict probability $p$. The same fit function is used for both graph-aware search and naive search.

#### 2.4 Scaling with compute

For graph-aware search, constructing the full conflict graph for $T$ candidate sequence pairs requires testing all pairwise combinations and therefore scales as

$$C_{\text{graph}}(T) = \frac{T(T-1)}{2} \approx \frac{T^2}{2}, \tag{37}$$

where one unit of $C_{\text{graph}}$ corresponds to the four NUPACK off-target calculations required to compare two sequence pairs.

Naive search has a more complicated cost structure because each candidate is tested only against the current independent set and is discarded as soon as the first conflict is detected.

Because naive search uses early rejection, each candidate is not necessarily compared with all sequence pairs in the current independent set. We can, however, compute the expected cost of comparing a new candidate against an existing independent set of size $I$.

We assign each comparison a probability of being executed. The first comparison ($j = 1$) is always executed and has probability 1. The second comparison ($j = 2$) is executed only if the first comparison has no conflict and has probability $1-p$. In general, the $j$-th comparison is executed only if the first $j-1$ comparisons have no conflict and therefore has probability $(1-p)^{j-1}$.

There are $I$ possible comparisons, each with its own assigned probability. Thus, we can compute the expected number of comparisons $C_{\text{cand}}$ as

$$C_{\text{cand}}(I) = \sum_{j=1}^{I} 1 \cdot (1-p)^{j-1} = \frac{1-(1-p)^I}{p}, \tag{38}$$

where we used the geometric-series formula again.

To estimate the expected comparison cost for adding one more sequence pair to the independent set, we use the earlier derived expected number of candidates $N(I)$ that need to be tested to increase the independent-set size from $I$ to $I+1$. If $N(I)$ candidates are tested on average, and each candidate has an expected comparison cost $C_{\text{cand}}(I)$, then the expected comparison cost for increasing the independent-set size by one is $N(I)C_{\text{cand}}(I)$.

Summing over all growth steps gives the expected cost to reach an independent set of size $I$:

$$C_{\text{naive}}(I) = \sum_{i=0}^{I-1} N(i)C_{\text{cand}}(i). \tag{39}$$

Using $N(i) = (1-p)^{-i}$ from equation 18, this becomes

$$C_{\text{naive}}(I) = \sum_{i=0}^{I-1} (1-p)^{-i}\frac{1-(1-p)^i}{p}. \tag{40}$$

This simplifies to

$$C_{\text{naive}}(I) = \frac{1}{p}\sum_{i=0}^{I-1} \left[(1-p)^{-i} - 1\right]. \tag{41}$$

Using $T(I) = \sum_{i=0}^{I-1}(1-p)^{-i}$ from equation 19, we obtain

$$C_{\text{naive}}(I) = \frac{T(I)-I}{p}. \tag{42}$$

Since $I$ grows only logarithmically with $T$, the leading large-$T$ scaling is

$$C_{\text{naive}}(T) \approx \frac{T}{p}. \tag{43}$$

Thus, with early rejection, the expected comparison cost of naive search grows approximately linearly and is a factor $1/p$ larger than the number of considered candidate sequence pairs. In contrast, full graph construction scales quadratically.

Lastly, we can combine the scaling estimates above to estimate how the independent-set size scales with computational cost for each algorithm. For naive search, the independent-set size scales at leading order as $I \approx \ln T/[-\ln(1-p)]$, while the computational cost scales as $C_{\text{naive}}(T) \approx T/p$. Thus, with $T \approx pC_{\text{naive}}$, we arrive at

$$I(C_{\text{naive}}) \approx \frac{\ln C_{\text{naive}}}{-\ln(1-p)}, \tag{44}$$

where we have omitted the constant offset $\ln p/[-\ln(1-p)]$.

For the graph-aware reference estimate, the independent-set size is expected to scale as $I \approx 2\ln T/[-\ln(1-p)]$, while full graph construction scales as $C_{\text{graph}}(T) \approx T^2/2$. Thus, with $T \approx \sqrt{2C_{\text{graph}}}$, we arrive at

$$I(C_{\text{graph}}) \approx \frac{\ln C_{\text{graph}}}{-\ln(1-p)}, \tag{45}$$

where we have omitted the constant offset $\ln 2/[-\ln(1-p)]$.

Thus, in this simplified scaling estimate, naive search and the graph-aware reference estimate show the same leading logarithmic scaling with computational cost. However, the graph-aware reference estimate does not describe the scaling of our graph-aware search implementation directly. Instead, it estimates the size of an independent set expected to exist in an idealized Erdős-Rényi conflict graph. Our graph-aware search implementation using the GMAX deletion heuristic is therefore expected to find independent sets that are smaller than this idealized reference value.

Thus, if conflict-graph construction is the limiting computational cost and the conflict graph is Erdős-Rényi-like, naive search is likely computationally favorable. In structured sequence-design conflict graphs, however, graph-aware search can still be advantageous because it uses global conflict information to enrich the independent set in globally low-conflict sequence pairs.

### 3 Conflict-probability analysis

To gain further insight into the benchmark results shown in Figure 4B of the main text, we selected the 12-mer benchmarks with and without a 5′ extension for additional conflict-graph analysis. We use this analysis to better understand the structure of the underlying conflict graphs and how this structure affects the behavior of naive search and hybrid search.

#### 3.1 General conflict structure

We first analyzed the initial graph-search subset used at the beginning of hybrid search to obtain a representative view of the conflict-graph structure. We used the hybrid-search runs with an initial graph-search subset containing $N = 900$ randomly sampled sequence pairs that had passed the energy pre-filters. For each of these sequence pairs, we calculated its conflict probability as the number of conflicts with the other $N - 1 = 899$ sequence pairs divided by $N - 1$. The resulting conflict-probability distributions are shown in Figure S1A. For the 5′ TTTT extension, the mean conflict probability was 0.148 with a standard deviation of 0.038. Without the extension, the mean conflict probability was 0.111 with a standard deviation of 0.036. For comparison, we considered an Erdős-Rényi graph, in which all conflict edges occur independently with the same probability. In this case, the number of conflicts $k_i$ of a sequence pair with the other $N - 1$ sequence pairs follows a binomial distribution with mean conflict probability $p$. The expected standard deviation of the measured per-sequence-pair conflict probability $p_i = k_i/(N - 1)$ is therefore $\sigma_{\mathrm{ER}} = \sqrt{p(1 - p)/(N - 1)}$. For the 5′ TTTT extension, this gives $\sigma_{\mathrm{ER}} = 0.0118$, compared with the measured value of 0.038. Without the extension, this gives $\sigma_{\mathrm{ER}} = 0.0105$, compared with the measured value of 0.036. The measured distributions are therefore more than threefold broader than expected for a graph with independent edges of uniform probability. This indicates that the conflict graph is not well described by an Erdős-Rényi graph. Instead, some sequence pairs are intrinsically more conflict-prone than others, producing a structured conflict graph with a broader distribution of conflict probabilities.

#### 3.2 Selection bias toward low-conflict sequence pairs

Next, we investigated how the conflict probabilities of the independent sets obtained by naive search and graph-aware search compare with randomly sampled sequence pairs outside the independent sets. For the 12-mer condition with a 5′ TTTT extension, graph-aware search found 49 sequence pairs from the initial graph-search subset with $N = 900$. We therefore selected these 49 sequence pairs and compared them with the first 49 sequence pairs found by naive search under the same condition. For the 12-mer condition without the 5′ extension, graph-aware search found 65 sequence pairs, so we compared these 65 sequence pairs with the first 65 sequence pairs found by naive search. For each condition, we then sampled 1000 additional sequence pairs that had passed the energy pre-filters and counted the number of conflicts between each sequence pair in the independent set and this sampled comparison pool. The resulting conflict-probability distributions are shown for naive search in Figure S1B and for graph-aware search in Figure S1C.

In all cases, the orthogonal sequence pairs had lower mean conflict probabilities than randomly sampled sequence pairs from the initial graph-search subset. Graph-aware search further reduced the mean conflict probability compared with naive search and produced a narrower distribution. For the $5'$ TTTT extension, the sequence pairs obtained by naive search had a mean conflict probability of 0.117 with a standard deviation of 0.041, whereas the sequence pairs obtained by graph-aware search had a mean conflict probability of 0.107 with a standard deviation of 0.024. Without the extension, the sequence pairs obtained by naive search had a mean conflict probability of 0.074 with a standard deviation of 0.032, whereas the sequence pairs obtained by graph-aware search had a mean conflict probability of 0.070 with a standard deviation of 0.026. These results indicate that both naive search and graph-aware search enrich the resulting independent set in globally low-conflict sequence pairs, with graph-aware search producing the slightly stronger enrichment.

This analysis also helps explain the relative performance of hybrid search and naive search in the long sequence benchmark. The advantage of hybrid search arises because the initial graph-aware search step enriches the starting independent set in sequence pairs with low conflict probability. These sequence pairs reject fewer candidates during the subsequent collection stage, making it easier to collect additional compatible sequence pairs. At the same time, the analysis also explains why naive search remains competitive. Although naive search does not use the full conflict-graph structure, its sequential rejection step still biases the growing independent set toward sequence pairs with relatively low conflict probability. Thus, both strategies enrich for globally low-conflict sequence pairs, but graph-aware search does so more directly and more strongly.

##### 3.3 Empirical scaling of independent-set size

Finally, we analyzed how the number of obtained orthogonal sequence pairs scales with the number of pre-filtered candidate sequence pairs considered during the search (Figure S1D). For graph-aware search, we included the initial graph-aware searches from the hybrid runs with subset sizes of 250, 450, and 900 sequence pairs, the separate graph-aware search condition containing 2235 sequence pairs, and the final graph-aware searches performed on the sequence pairs obtained during the collection stage. For comparison, we plotted the naive-search trajectory on the same axis, where the x-axis likewise corresponds to the number of candidate sequence pairs that passed the energy pre-filters. This representation allows the growth of the independent set to be compared between naive search and the two graph-aware search stages of hybrid search using the same measure of candidate-pool usage. The analysis was performed separately for the 12-mer conditions with and without a $5'$ extension.

To compare the scaling behavior, we fitted the function $A \ln(1 + x/B)$ to the naive-search trajectory and to the combined graph-aware-search data from the initial and final graph-aware searches. In both extension conditions, the initial and final graph-aware-search data followed the same trend and could be fitted together as a single dataset. This indicates that the graph-aware search step behaves similarly whether the candidate sequence pairs are obtained by random sampling or by collection after compatibility testing against the initial independent set.

The logarithmic fit captured the trajectories of both naive search and graph-aware search. Without a $5'$ extension, the fitted prefactor was $A = 10.10$ for naive search and $A = 21.40$ for graph-aware search. With the $5'$ TTTT extension, the fitted prefactor was $A = 5.88$ for naive search and $A = 12.61$ for graph-aware search. Thus, graph-aware search showed a larger fitted logarithmic prefactor in both cases, consistent with the scaling estimate above. Overall, graph-aware search outperformed naive search when compared at the same number of pre-filtered candidate sequence pairs considered.

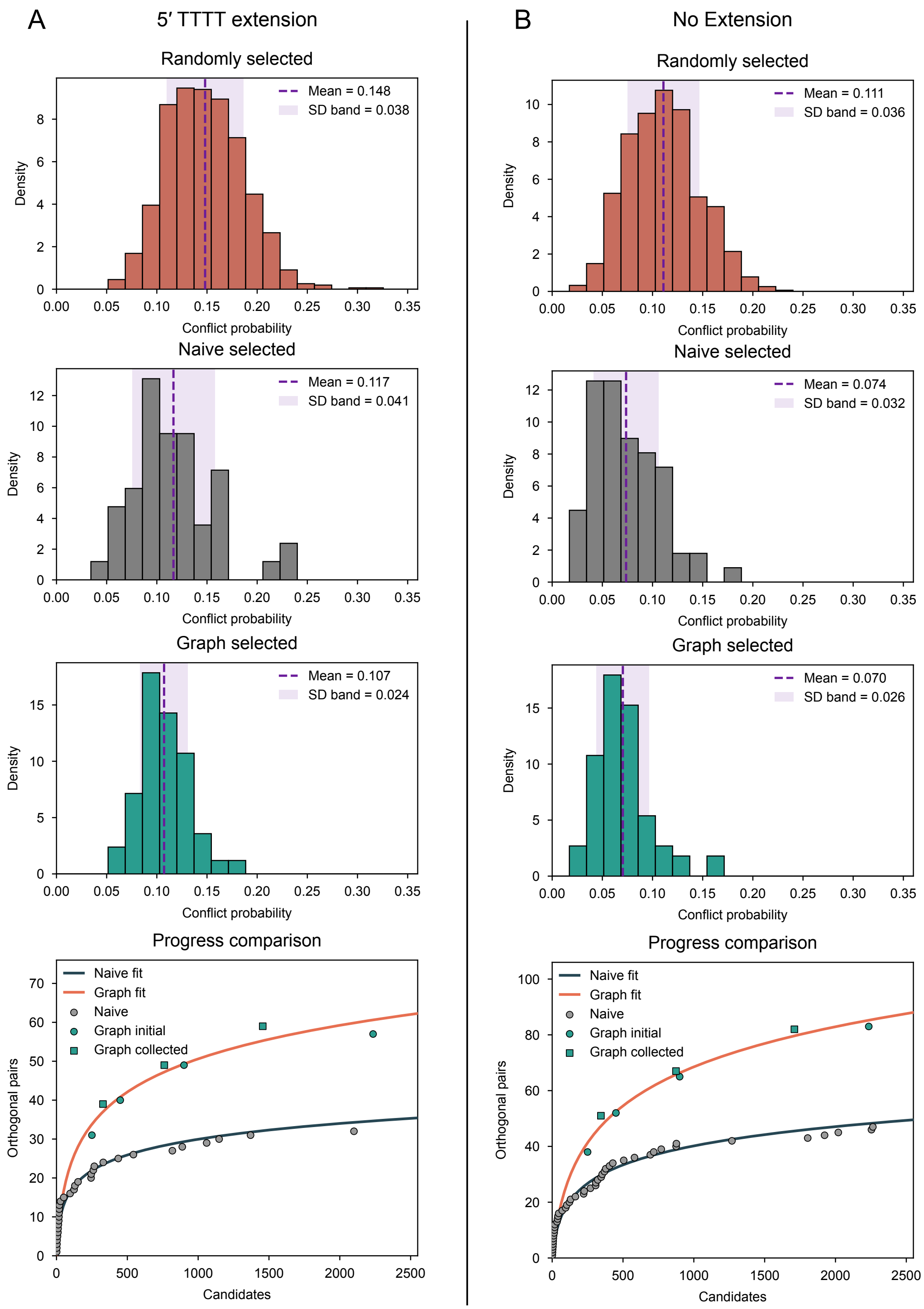

**Supplementary Figure S1: Conflict-probability analysis of representative 12-mer sequence-pair graphs.** All panels correspond to the 12-mer benchmark shown in Figure 4B of the main text. Left panels show sequence pairs with a 5′ TTTT extension, and right panels show sequence pairs without a 5′ extension. **(A)** Conflict-probability distributions for the randomly selected initial graph-search subsets. **(B)** Conflict-probability distributions of sequence pairs obtained by naive search, measured against 1000 randomly sampled sequence pairs outside the independent set. **(C)** Conflict-probability distributions of sequence pairs obtained by graph-aware search, measured against 1000 randomly sampled sequence pairs outside the independent set. **(D)** Number of orthogonal sequence pairs obtained as a function of the number of considered pre-filtered candidate sequence pairs for naive search and graph-aware search. Graph-aware-search data include the initial graph-aware searches with subset sizes of 250, 450, 900, and 2235 sequence pairs and the final graph-aware searches from the hybrid runs. Logarithmic fits are shown as lines.

### 4 Perturbation fraction analysis

The perturbation fraction controls how strongly the current independent set is modified during each iterative perturbation step. In our implementation, it specifies the fraction of vertices in the full graph that are temporarily reintroduced into the current independent set before the resulting conflicts are repaired with the GMAX deletion heuristic and the subsequent cleanup step. Small perturbation fractions make only minor changes to the current independent set and may therefore limit exploration of alternative independent sets, whereas very large perturbation fractions strongly disrupt the current independent set and may reduce the benefit of starting from an already good one.

To choose a single perturbation fraction for benchmarking, we tested a range of values across multiple representative conditions. We used the same pre-filtering conditions as in the benchmarks shown in Figure 4 of the main text and considered core binding-domain lengths of 10, 16, and 20. For each length, we tested initial graph-search subset sizes of 250, 450, and 900 sequence pairs. For every combination of length and initial subset size, we evaluated perturbation fractions of 0.025, 0.05, 0.10, 0.15, 0.20, 0.25, 0.30, 0.40, 0.50, 0.60, 0.70, 0.80, 0.90, 0.95, and 1.00. Each condition was repeated with three random seeds.

The results are shown in Figure S2. Each panel corresponds to one core binding-domain length and contains the results for the three tested initial subset sizes. The dashed lines show the average independent-set size after the initial graph-aware search heuristic, before iterative perturbation, averaged over the three random seeds. The solid lines with error bars show the corresponding independent-set size after 1000 perturbation iterations. Across all tested conditions, very small and very large perturbation fractions performed worse than intermediate values, with the best results often obtained for perturbation fractions near 0.5. We used a perturbation fraction of 0.2 for the benchmark calculations. This value was initially chosen based on the $L = 10$ condition and was slightly below the best-performing range in the full parameter sweep. However, the qualitative benchmark conclusions do not depend on choosing the optimal perturbation fraction.

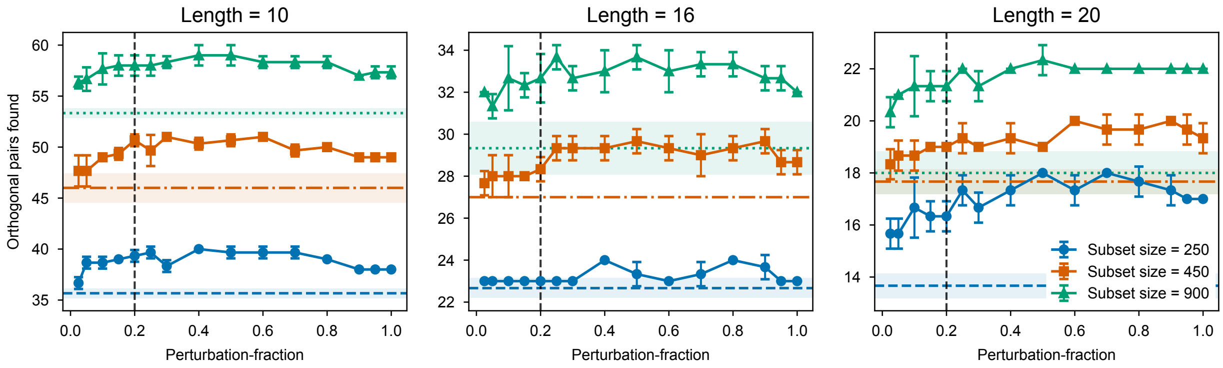

**Supplementary Figure S2: Perturbation-fraction analysis for graph-aware search.** Independent-set sizes are plotted against the perturbation fraction for core binding-domain lengths of 10, 16, and 20, using the same sequence-layout constraints and thermodynamic pre-filtering criteria as in the benchmarks shown in Figure 3 of the main text. Within each panel, data are shown for initial graph-search subset sizes of 250, 450, and 900 sequence pairs. Dashed lines indicate the average independent-set size after the initial graph-aware search heuristic, before iterative perturbation, averaged over three random seeds. The shaded regions around the dashed lines indicate the corresponding standard deviations. Solid lines with data points indicate the corresponding independent-set sizes after 1000 perturbation iterations, and error bars indicate standard deviations. The vertical line marks the perturbation fraction of 0.2 used for the benchmarks shown in Figure 4 of the main text.

### 5 Long sequence benchmark at 25 °C

We repeated the long sequence benchmark at 25 °C. The benchmark used the same sequence-layout constraints, search strategies, initial graph-search subset sizes, and fixed NUPACK-call budget as the long-sequence benchmark shown in Figure 4B of the main text.

Energy limits were recalculated for 25 °C. The allowed on-target energy range was chosen separately for each core binding-domain length and extension condition using the same standard-deviation-based strategy as in the main benchmark. The off-target limit was chosen from the same bound-fraction criterion used for the long-sequence benchmark, giving $-7.845$ kcal/mol at 1 $\mu$M strand concentration. The self-folding energy limit was chosen from the same minimum unpaired-fraction criterion, giving $-0.954$ kcal/mol.

The results are shown in Figure S3. This benchmark tests whether the relative performance of naive search and hybrid search is preserved when the thermodynamic energy limits are recalculated for a lower experimental temperature.

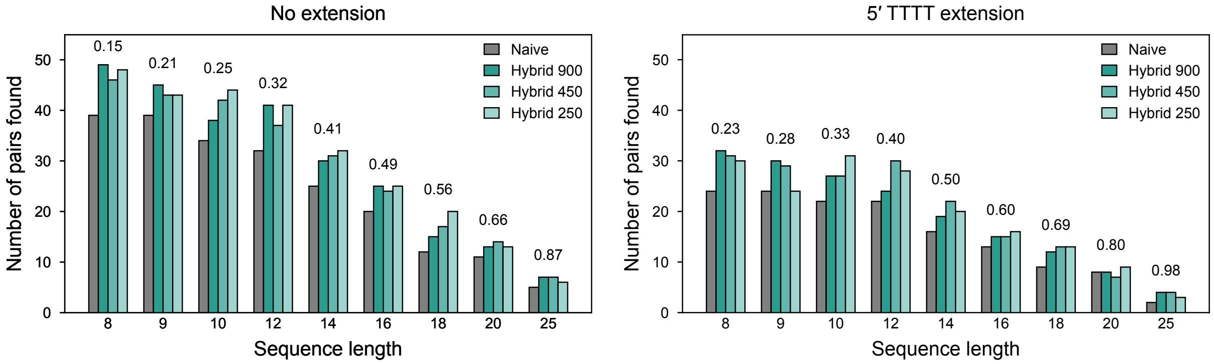

**Supplementary Figure S3: Long sequence benchmark at 25 °C.** Number of orthogonal sequence pairs found by naive search and hybrid search for core binding domains with and without a 5′ TTTT extension. Naive search and hybrid search were run with a fixed computational budget of $10^7$ NUPACK calls. For hybrid search, initial graph-search subset sizes of 250, 450, and 900 sequence pairs were tested. The perturbation fraction was set to 0.2, and graph-aware search was run for 1000 iterations. Sequence pairs containing GGGG were removed to avoid G-quadruplex-forming sequences. Energy limits were recalculated for 25 °C. The allowed on-target association free-energy range was chosen separately for each core binding-domain length and extension condition using the same standard-deviation-based strategy as in the main benchmark. The off-target energy limit was set to $-7.845$ kcal/mol, and the self-folding energy limit was set to $-0.954$ kcal/mol.

### 6 Supplementary Methods

#### 6.1 Graph-aware search algorithm

**Vertex cover and independent set.** We implemented the graph-aware search algorithm from the perspective of a vertex cover, which is complementary to an independent set. A vertex cover is a set of vertices that touches every edge in the graph. In other words, each edge has at least one endpoint in the vertex cover. The vertices that are not in the vertex cover automatically form an independent set: if two vertices outside the vertex cover were connected by an edge, that edge would not be covered and the vertex cover would not be complete. Thus, searching for a large independent set is equivalent to searching for a small vertex cover.

**Basic algorithm.** The basic removal strategy is known as the GMAX maximum-degree deletion heuristic (*6*). In this strategy, each vertex in the conflict graph is labeled by its number of edges, corresponding to its degree in the graph. The vertex with the highest number of edges is removed first and added to the vertex cover. The number of conflicts is then recalculated for the remaining vertices, and the procedure is repeated. Once no edges remain, the collected vertices form a vertex cover, and the remaining vertices form an independent set.

**Self-loops.** A self-loop is an edge that connects a vertex to itself. In the conflict graph, self-loops occur when at least one strand of a sequence pair forms a homodimer stronger than the chosen off-target limit. Before applying the GMAX deletion heuristic, vertices with self-loops are removed first and added to the vertex cover. Homodimer-forming sequence pairs are usually removed during the energy pre-filtering stage. However, for the full conflict graphs used in Figure 4A of the main text, self-loops were retained during graph construction and eliminated before running the deletion heuristic.

**Tie-breaking.** During the GMAX deletion heuristic, several vertices can have the same highest number of conflict edges. We therefore use a tie-breaking rule based on the overlap between high-conflict vertices. Among the vertices with the highest number of edges, we remove the vertex with the smallest overlap count. The overlap count is defined as the number of neighboring vertices that also have the highest number of edges. The rationale is that a vertex with a low overlap count is less likely to be resolved indirectly by removing another high-conflict vertex in a following step. In contrast, vertices with high overlap counts share many conflicts with other high-conflict vertices, so their number of edges may decrease during subsequent removal steps anyway. If this rule results in another tie, one of the remaining tied vertices is picked at random.

**Cleanup.** After running the GMAX deletion heuristic, we apply a cleanup routine to identify vertices that were excluded unnecessarily. During iterative vertex removal, a vertex may be removed early, even though all vertices in conflict with it are removed later. In this case, the final independent set can be expanded by reintroducing such excluded vertices. We therefore check all excluded vertices one by one and add a sequence pair back to the independent set if its addition does not create any conflict with the current independent set. When several excluded vertices are admissible, they are reconsidered one by one.

**Iterative perturbation.** Starting from an existing independent set, we randomly reintroduce a subset of previously excluded vertices. This perturbation generally reintroduces conflicts. We therefore repair the perturbed set by applying the same GMAX deletion heuristic to the resulting conflict graph. After this repair step, we again apply the cleanup step to all vertices in the full graph. If the resulting independent set is larger than the previously best set, it is retained and used as the starting point for the next perturbation iteration. This procedure allows vertices in the independent set to be exchanged for previously excluded vertices. In particular, vertices that block the addition of several other vertices can be removed during the repair step, allowing a larger independent set to be found. This strategy extends the general idea of local-search improvement for maximum independent set problems (*7*). This introduces an additional parameter, the perturbation fraction, which specifies the fraction of all vertices that is temporarily reintroduced into the independent set during each perturbation step. The choice of this parameter is analyzed in Section 4 and Figure S2. In our implementation, the perturbation is carried out equivalently from the complementary vertex-cover perspective by temporarily removing a random subset of vertices from the current vertex cover and then repairing the resulting uncovered edges with the GMAX deletion heuristic.

#### 6.2 Sequence sampling

**Sequence-space size.** For a core binding-domain length $L$, the number of possible sequence pairs scales as $4^L/2$, because each core sequence is paired with its reverse complement and the two orientations define the same sequence pair. For larger core binding domains, it is therefore impractical to generate and store all possible sequence pairs before the search begins. At the same time, the graph-aware algorithms require each sequence pair to be uniquely identifiable. We therefore assign each sequence pair a persistent unique ID and use a sequence generator that creates sequence pairs on demand. The prefiltered pool shown schematically in Figure 2 of the main text is not constructed explicitly, but is defined implicitly by the sequence-layout constraints and energy criteria.

**Sampling.** The sequence generator is initialized with the core binding-domain length, optional $5'$ and $3'$ extensions, excluded sequence motifs, and the region to which motif exclusion is applied. To sample a sequence pair, a random core sequence is generated, paired with its reverse complement, and extended with the specified $5'$ and $3'$ sequences. Sequence pairs containing excluded motifs are discarded. The two strands are then sorted into a canonical order and checked against an internal register of previously sampled sequence pairs. The generator returns the corresponding persistent ID, assigning a new ID only if the sequence pair has not been sampled before.

#### 6.3 Sequence pre-filtering

**Pool exhaustion.** Naive search, the collection stage of hybrid search, and the generation of the initial graph-search subset all require a continuous supply of previously unseen sequence-pair IDs. Because the sequence generator can return sequence pairs that were sampled before, each algorithm maintains a list of already seen IDs and discards repeated IDs. For short core binding domains, the accessible sequence-pair pool can become exhausted, making it increasingly unlikely to sample a new ID. We therefore terminate sampling after $M = 10^6$ consecutive attempts fail to produce a previously unseen sequence-pair ID. This stopping rule can be interpreted probabilistically. If a fraction $p$ of the accessible sequence-pair pool remains unseen, the probability of failing to sample a new ID in $M$ consecutive attempts is $P_M = (1 - p)^M$, giving $p = 1 - P_M^{1/M}$. For $M = 10^6$ and $P_M = 0.01$, this gives $p \approx 4.6 \times 10^{-6}$. Thus, after $10^6$ consecutive failed attempts, there is 99 % confidence that less than approximately $4.6 \times 10^{-6}$ of the accessible sequence-pair pool remains unseen, assuming random sampling from the accessible pool.

**Energy pre-filtering.** Each newly sampled sequence pair is first evaluated using the thermodynamic criteria that depend only on intrinsic properties of the sequence pair. First, the on-target association free energy is computed together with the self-folding free energies of both individual strands. The sequence pair is rejected if the on-target energy does not lie within the selected allowed range, or if at least one of the two strands has a self-folding free energy below the selected limit. Sequence pairs that pass these criteria are then tested for same-strand homodimer formation. If either strand forms a homodimer stronger than the off-target limit, the sequence pair is rejected. Homodimerization is formally an off-target interaction. However, we apply this filter at the pre-filtering stage to prevent sequence pairs that are already unsuitable from entering the more expensive downstream computations. Equivalently, such sequence pairs would be removed during graph-aware search by the initial elimination of vertices with self-loops.

#### 6.4 Naive search

In naive search, sequence pairs are generated on demand as described in Section 6.2 and subjected to the same energy pre-filtering described in Section 6.3. Candidate sequence pairs that pass these filters are tested against the sequence pairs already retained in the growing orthogonal set. For each candidate-retained pair comparison, all four possible off-target interactions are evaluated. The candidate sequence pair is accepted only if none of these interactions is stronger than the chosen off-target limit. To reduce computation time, each candidate is discarded immediately after the first off-target-limit violation is detected. The search continues until the available sequence space is exhausted, a predefined NUPACK call budget is reached, or the search is stopped manually.

#### 6.5 Hybrid search

**Initial graph-aware search.** Hybrid search uses the same sequence sampling and energy pre-filtering procedures as described in Sections 6.2 and 6.3. First, sequence pairs are sampled until a predefined initial subset size is reached. All off-target interactions between sequence pairs in this subset are then computed, and the complete conflict graph is constructed from these energy values. The graph-aware search described in Section 6.1 is then applied to obtain an initial independent set.

**Collection stage.** In the second stage, additional sequence pairs are sampled and energy pre-filtered. Each candidate that passes the pre-filters is tested against all sequence pairs in the initial independent set. For each comparison between a candidate sequence pair and a sequence pair from the initial independent set, all four possible off-target interactions are evaluated. The candidate is added to a collection pool only if none of these interactions are stronger than the chosen off-target limit. As in naive search, candidates are discarded immediately after the first off-target-limit violation is detected to reduce computation time. The collection pool is therefore compatible with the initial independent set by construction, but is not necessarily internally conflict-free because collected candidates are not tested against each other during this stage.

**Final graph-aware search.** After candidate collection, a second conflict graph is constructed only for the collected sequence pairs. To construct this graph, all off-target interactions between collected sequence pairs are computed, and edges are added for sequence-pair combinations that violate the chosen off-target limit. Graph-aware search is then applied to this graph to identify an additional independent set. Because all collected sequence pairs have already been tested against the initial independent set, this additional independent set can be combined directly with the initial independent set to obtain the final orthogonal sequence-pair set.

**Stopping criteria.** Hybrid search stops when the sequence pool is exhausted, when a predefined NUPACK call budget is reached, or when the search is stopped manually. During the collection stage, the search also reserves sufficient NUPACK calls for the final graph-aware search on the collected sequence pairs. For a collection pool containing $N$ sequence pairs, the final graph construction requires evaluating four off-target interactions for each of the $N(N-1)/2$ sequence-pair combinations. The required number of NUPACK calls is therefore estimated as $4 \cdot N(N-1)/2 = 2N(N-1)$. Candidate collection is stopped before this estimated final graph-construction cost would cause the total NUPACK call budget to be exceeded.

#### 6.6 Benchmarking

**General benchmark setup.** All benchmarks were performed using the thermodynamic conditions shown in Figure 1B of the main text. Unless stated otherwise, NUPACK calculations were carried out for DNA at 37 °C, with 50 mM NaCl and 25 mM $MgCl_2$. Sequence pairs containing GGGG (and CCCC) were removed to avoid G-quadruplex-forming sequences. We benchmarked core binding-domain lengths from 4 to 25 nucleotides and compared sequence pairs with and without a $5'$ TTTT extension. The benchmark was separated into short-sequence and long-sequence regimes. For short core binding domains with $L \leq 7$, all off-target interactions could be precomputed, allowing direct comparison with graph-aware search. For longer core binding domains with $L > 7$, full graph construction from the complete sequence space became impractical, so the search strategies were compared under a fixed NUPACK-call budget. In addition to naive search and hybrid search, we included a graph-aware search condition in which the full computational budget was allocated to constructing and searching a single conflict graph.

**Energy limits.** The allowed on-target energy range was chosen separately for each core binding-domain length and extension condition. For the short-sequence benchmark, the full candidate pool was generated for each condition, and the on-target energy distribution was computed over all candidate sequence pairs. For the long-sequence benchmark, full pool enumeration was not feasible, so the on-target energy distribution was estimated from 2000 sampled sequence pairs for each condition. In both cases, the allowed on-target range was defined from the mean of the corresponding distribution to one standard deviation below the mean.

This choice keeps the selected on-target binding strength comparable across sequence lengths and extension conditions while excluding the strongest-binding tail of the distribution. Stronger on-target interactions also tend to be associated with stronger off-target interactions, so excluding this tail reduces the number of sequence pairs that pass the on-target filter but fail later off-target filtering.

For all benchmarks, the self-folding energy limit was chosen such that the predicted unpaired fraction remained above 0.2. At 37 °C, this corresponds to $-0.99$ kcal/mol. Off-target limits were chosen differently for the short sequence and long sequence benchmarks, as described below.

**Short sequence benchmark.** For the short sequence benchmark, the full candidate pool was generated for each length and extension condition. On-target and self-folding energies were computed for all sequence pairs, and sequence pairs within the allowed on-target range were retained for off-target matrix construction.

All off-target interactions between retained sequence pairs were then precomputed and stored as cached energy matrices. These matrices were used for naive search, hybrid search, and graph-aware search, so that all strategies were compared on the same sequence pool and the same off-target energy data. Implementation details of the offline benchmark algorithms are provided below.

For short core binding domains, a fixed bound-fraction off-target limit is not a useful benchmark parameter because binding energies depend strongly on core length. An energy limit that is meaningful for 7-mer cores would be too permissive for 4-mer cores, whereas a limit suitable for 4-mer cores would be too strict for 7-mer cores. We therefore chose off-target limits by targeting fixed mean conflict probabilities for each length and extension condition. Conflict probability was defined as the number of conflict edges divided by the number of possible edges, including self-loops. Target mean conflict probabilities of 0.1, 0.2, and 0.3 were used, and the corresponding off-target limits are listed in Supplementary Table S1. For hybrid search, initial subset sizes of 250, 450, and 900 sequence pairs were tested. The perturbation fraction was set to 0.2, and graph-aware search was run for 1000 iterations.

**Offline benchmark algorithms.** We created offline versions of naive search and hybrid search that operate on precomputed off-target energy values, allowing them to be compared directly with the full graph-aware search, which works exclusively on precomputed off-target energy data. In all offline searches, the on-target filter is not applied during the search itself because the precomputed candidate pool already contains only sequence pairs within the allowed on-target energy range.

The offline naive search iterates through the candidate pool in a shuffled order and accepts a sequence pair only if it satisfies the self-folding and homodimer filters and has no conflict with the already accepted sequence pairs, analogous to the on-demand naive search described in Section 6.4. The required energy values are read from the precomputed dataset rather than recalculated during the search.

The offline hybrid search follows the same initial graph-aware search, collection, and final graph-aware search structure described in Section 6.5, but uses cached energy matrices instead of recalculating NUPACK energies. As in the offline naive search, candidate sequence pairs are required to satisfy the self-folding and homodimer filters before they enter the initial graph-search subset or the collection pool. Initial graph-search subset sizes of 250, 450, and 900 sequence pairs were tested. The perturbation fraction was set to 0.2, and the graph-aware search procedure was run with 1000 iterations.

The offline graph-aware search constructs the conflict graph from the cached matrices after applying the self-folding filter and then applies the graph-aware search described in Section 6.1. Homodimer-forming sequence pairs are not removed by a separate pre-filter in the graph-aware search. Instead, they appear as vertices with self-loops and are removed during the initial self-loop elimination step of the graph-aware search. The perturbation fraction was set to 0.2, and the search procedure was run for 1000 iterations.

**Long sequence benchmark.** For the long sequence benchmark, full off-target matrix construction for the complete sequence space was not feasible. We therefore compared naive search, hybrid search, and graph-aware search under a fixed computational budget of $10^7$ NUPACK calls. Naive search and hybrid search used the on-demand implementations described in Sections 6.4 and 6.5. For hybrid search, initial subset sizes of 250, 450, and 900 sequence pairs were tested. For the graph-aware search condition, the full computational budget was allocated to the initial graph construction and search, resulting in a graph containing approximately 2235 sequence pairs. The off-target limit was set by requiring off-target interactions to remain below $1\,\%$ bound at $1\,\mu$M strand concentration, corresponding to $-8.16\,$kcal/mol. The perturbation fraction was set to 0.2, and graph-aware search was run for 1000 iterations.

#### 6.7 Graphical user interface

The graphical user interface was implemented in Streamlit and is intended as the main user-facing interface for applying the sequence-selection algorithms. The application is organized into five workflow tabs: Selection Helper, Pilot Analysis, Off-Target Limit, Sequence Search, and Load Results. The Selection Helper tab supports conversion between bound-fraction or unpaired-fraction criteria and the corresponding energy limits. The Pilot Analysis and Off-Target Limit tab allow the selection of energy limits. User-selected energy limits are displayed as vertical lines in the corresponding plots and update interactively when the selected values are changed. The Sequence Search tab runs the sequence-selection algorithms using the selected parameters. Search progress and intermediate computations are reported through an integrated console-like log window that refreshes automatically while a search or analysis task is running. When a search is finalized, the selected sequence-pair library and associated parameters are saved as an Excel report. The Load Results tab allows previously generated reports to be loaded and visualized.

### 7 Supplementary Tables

**Supplementary Table S1: Off-target limits used for the short-sequence benchmark in Figure 4 of the main text.**

| Core length $L$ | Target conflict probability | Off-target limit (kcal/mol) | |
|---|---|---|---|
| | | No 5′ extension | 5′ TTTT extension |
| 4 | 0.1 | -5.143 | -6.218 |
| 4 | 0.2 | -4.828 | -5.842 |
| 4 | 0.3 | -4.484 | -5.376 |
| 5 | 0.1 | -6.141 | -6.877 |
| 5 | 0.2 | -5.643 | -6.329 |
| 5 | 0.3 | -5.347 | -6.025 |
| 6 | 0.1 | -6.583 | -7.253 |
| 6 | 0.2 | -6.086 | -6.634 |
| 6 | 0.3 | -5.751 | -6.342 |
| 7 | 0.1 | -7.050 | -7.660 |
| 7 | 0.2 | -6.511 | -6.995 |
| 7 | 0.3 | -6.199 | -6.620 |
